# A complex interplay between balancing selection and introgression maintains a genus-wide alternative life history strategy

**DOI:** 10.1101/2021.05.20.445023

**Authors:** Kalle Tunström, Alyssa Woronik, Joseph J. Hanly, Pasi Rastas, Anton Chichvarkhin, Andrew D Warren, Akito Kawahara, Sean D. Schoville, Vincent Ficarrotta, Adam H. Porter, Ward B. Watt, Arnaud Martin, Christopher W. Wheat

## Abstract

Alternative life-history strategies (ALHS) are genetic polymorphisms generating phenotypes differing in life histories that generally arise due to metabolic resource allocation tradeoffs. Althouigh ALHS are often be limited to a single sex or populations of a species, they can, in rare cases, be found among several species across a genus. In the butterfly genus *Colias*, at least a third of the species have a female limited ALHS called Alba. While many females develop brightly pigmented wings, Alba females reallocate nitrogen resources used in pigment synthesis to reproductive development, producing white-winged, more fecund females. Whether this ALHS evolved once or many times, and whether it has moved among species via introgression or been maintained via long-term balancing selection, has not been established. Answering these questions presents an opportunity to investigate the genetic basis and evolutionary forces acting upon ALHS, which have rarely been studied at a genus level. Here we identify the genetic locus of *Alba* in a second *Colias* species, allowing us to compare this with previous results in a larger phylogenetic context. Our findings suggest *Alba* has a singular origin and has been maintained in *Colias* through a combination of balancing selection and introgression for nearly one million years and at least as many generations. Finally, using CRISPR/Cas9 deletions in the cis-regulatory region of the *Alba* allele, we demonstrate that the *Alba* allele is a modular enhancer for the *BarH1* gene and is necessary for the induction of the ALHS, which potentially facilitates its long-term persistence in the genus.

## Introduction

Species vary in their life histories, allocating resources in differing amounts to growth, maintenance, and reproduction in an attempt to maximize fitness (Rose and Mueller, 1993). Within species, individuals also vary, whether plastically or genetically, in how they allocate resources. When genetically determined and causing distinctly different phenotypes, such polymorphisms within species are called alternative life-history strategies (ALHS)(Gross, 1996). Examining why some species remain polymorphic, as opposed to becoming fixed for one strategy, will provide insight into the ecological and evolutionary forces that shape ALHS during the adaptive diversification of populations and species (Jamie and Meier, 2020), and in turn, will inform our understanding of how complex traits evolve (Zakas et al., 2018).

Alternative life-history strategies are unstable by nature, with a single phenotype predicted to fixate within species over time (Ford, 1945; Llaurens et al., 2017). Thus, when an ALHS is observed in multiple species within a genus, at least one of the following must be true: **a)** the ALHS arose once and has been maintained by some form of balancing selection (Mérot et al., 2020a), **b)** the ALHS evolved independently via novel mutations causing similar phenotypes in separate species (Blow et al., 2021; Yassin et al., 2016), or **c)** the ALHS moved between species via introgression (Dasmahapatra et al., 2012). Distinguishing the relative role these nonexclusive alternatives have played in the evolution of an ALHS is challenging, especially as the genetic basis of ALHS is poorly understood in most species, let alone among many related species. Thus, an understanding of how ALHS evolve remains elusive.

Although butterflies in the genus *Colias* are characterized by their yellow to orange wing coloration, a third of the approximately 90 butterfly species of this genus have a female limited ALHS called Alba, where female wings are white (Limeri and Morehouse, 2016; Remington, 1954). The remaining species appear to be fixed for Alba or colored wings. Alba females reallocate metabolic resources from wing pigmentation to reproductive development, resulting in white wings, rather than the colored wings shared by males and the remaining females (Descimon and Pennetier, 1989; Graham et al., 1980; Nielsen and Watt, 1998; Watt, 1973; Woronik et al., 2018). Genetic studies in six *Colias* species consistently found that Alba is caused by a single dominant, autosomal locus (Remington, 1954). In *Colias crocea*, a Eurasian species polymorphic for Alba, the locus of the ALHS has been mapped to a transposable element insertion downstream of the gene *BarH1* (Woronik et al., 2019). This insertion leads to a gain of function of the *BarH1* gene in the developing wings, resulting in a lack of pigment granules in Alba wing scales (Woronik et al., 2019), in addition to a previously described reduction of pterin pigments (Watt 1973). In contrast, white color in other Pieridae species is primarily due to white pteridine pigment granules (Giraldo and Stavenga, 2007; Watt and Bowden, 1966; Wijnen et al., 2007).

This detailed knowledge of the genetic basis of this ALHS presents an opportunity to investigate the evolutionary dynamics of a complex life-history trait among species. The Alba life-history tradeoff between color production and reproductive investment has been studied in detail in both North American *C. eurytheme* and Eurasian *C. crocea* (Descimon and Pennetier, 1989; Woronik et al., 2018). Although Alba in both species appears to be a similar ALHS, the genetic basis of the Alba phenotype in any North American species is unknown. Since these two species likely last shared an ancestor before the North American and Eurasian clades separated, and these two clades contain the vast majority of *Colias* species, resolving whether these two species have the same or independent Alba phenotypes is necessary in order to understand the prevalence of this ALHS across the genus. While balancing selection can maintain life history polymorphisms, there is very little evidence that such a mechanism alone can maintain ALHS over deep evolutionary time. Shared ancestry, however, does not necessarily require that a polymorphism be ancient. Among *Heliconius* butterflies, regulatory units affecting wing color patterning readily move among populations and species via introgression, followed by strong directional selection leading to fixation (Morris et al., 2020; Wallbank et al., 2016; Westerman et al., 2018a). Whether introgression could play a similar role for an ALHS like Alba is unknown.

Here we set out to test among the alternative (single or multiple origins) and complimentary (balancing selection, introgression) evolutionary mechanisms responsible for the prevalence of the Alba ALHS (being fixed, absent or polymorphic) across the *Colias* genus. Using a combination of phylogenetic analyses, GWAS, and genetic manipulation, we find evidence that 1) the Alba polymorphism arose once, likely at the root of the genus, 2) it is maintained by balancing selection with introgression, and 3) that the *Alba* allele acts as a modular enhancer controlling this trans-specific ALHS.

## Results

### Phylogenomic analysis

A chromosome-level genome for *C. eurytheme* was generated and used as a reference for aligning whole-genome sequence data from 21 species representing the global distribution of *Colias* and diverse Alba phenotypes (Fig. 1; Supplementary Table 1). After haplomerging and polishing, the final haploid genome was 328 MB, with an N50 of 5.2MB across 108 scaffolds and high gene completeness with low duplication (97.7% of expected single-copy genes were complete and unique, i.e., 5166 of 5286 BUSCO genes; Supplementary Table 2). We then generated and used a linkage map to assemble chromosomes, finding evidence for 31, which is the expected number for most *Colias* (Maeki and Remington, 1960). The *C. eurytheme* chromosomal structure was highly syntenic with the standard organization of Lepidoptera chromosomes (Supplementary Fig. 1). Annotation of the resulting chromosome level assembly identified 18077 transcripts from 16842 genes.

**Figure 1.**
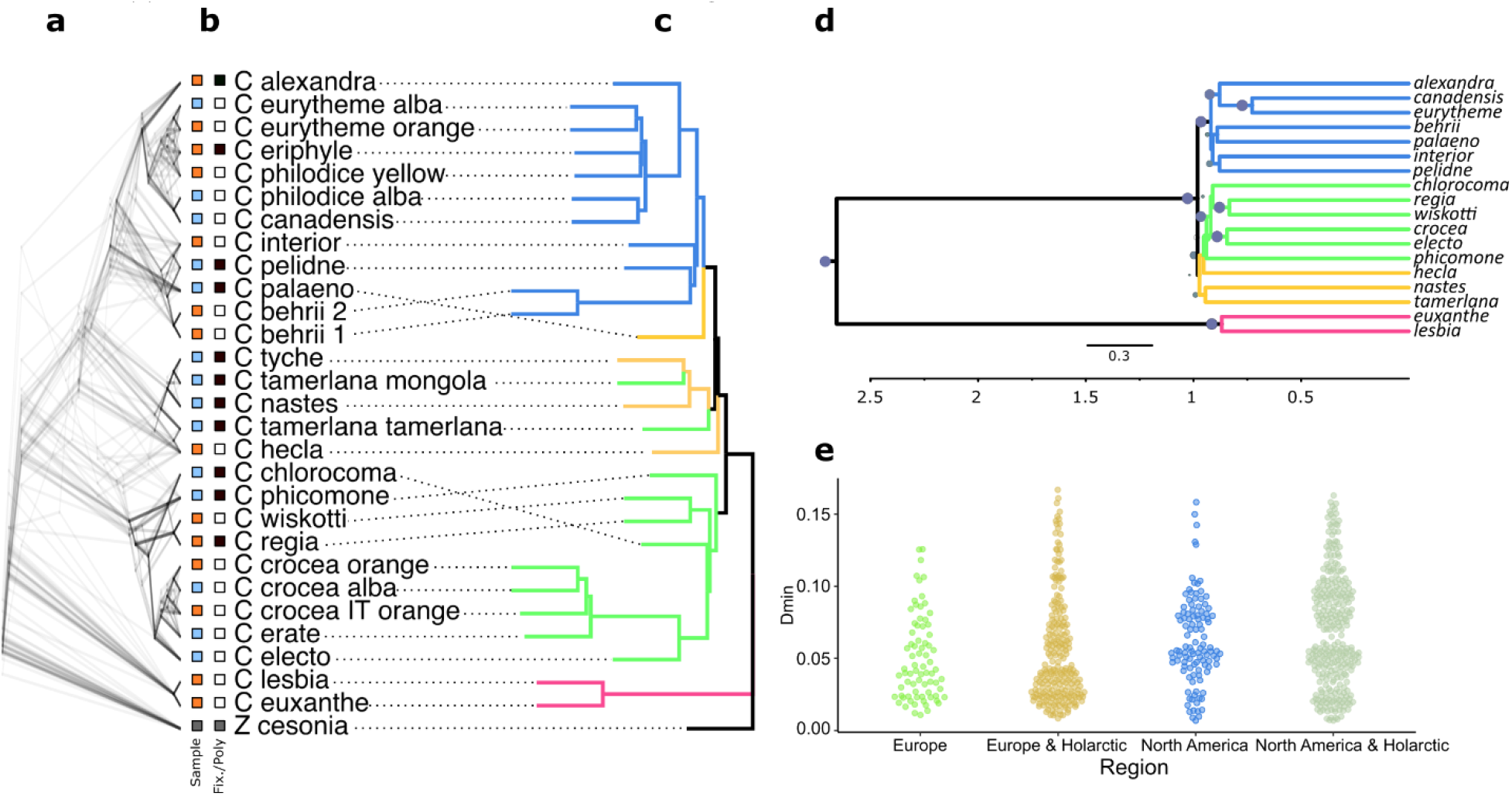
Evolutionary relationships among *Colias* butterfly species, their distribution, Alba phenotypes, and introgression. **a.** Thirty-one species trees are shown overlapping, one from each chromosome generated using gene trees based upon a single exon per gene (on average 129 genes / chromosome; n=4011 genes). **b.** For each specimen, first column indicates wing color morph (blue = Alba, orange = colored), while second column Alba morph type (black boxes = species is fixed for one morph, white = species is polymorphic). **c**. Species tree generated from all BUSCO exon trees, with branches color-coded by their samples regional distribution (Blue = North America, Orange = Holarctic, Green=Eurasia and Africa, Purple = South America). d. SNAPP tree generated using 1000 SNPs with secondary calibration, with millions of years on X axis. Dots upon nodes indicate posterior support, with values > 0.92 indicate by large blue circles. **e.** Distribution of minimal D-statistic of all species-trios that showed significant levels of introgression (BH corrected p < 0.05). Trios are grouped by Eurasian or North American regions, and then combined with the Holarctic species.

We mined these alignments for single-copy orthologs in Lepidoptera, then used the longest exon per gene to estimate maximum-likelihood gene trees, followed by species tree estimation using ASTRAL. Although there was extensive conflict among gene trees (Fig 1a; Supplementary Fig. 2; (n=4,244)), the species tree (Fig. 1c) supports three conclusions: species from South America are sister to the rest of the *Colias* genus, the major divergence within *Colias* is between a North American and a Eurasian + African clade, and circumpolar taxa, which we refer to as Holarctic, fall between and among these two major clades (Fig. 1c). In order to further assess these relationships in a multispecies coalescent framework, and to estimate divergence time among the North American and the Eurasian + African clades, we used a Bayesian species-tree inference approach (SNAPP) calibrated on an age estimate of when *Colias* and its sister genus *Zerene* last shared a common ancestor (Chazot et al., 2019). SNAPP is computationally demanding, necessitating a down sampling of data and taxa. Following recommendations (Stange et al., 2018), we analyzed a random set of 1000 SNPs selected from a reduced set of taxa, selected to remove redundancy among closely related species while retaining regional diversity. The SNAPP estimated phylogeny was largely concordant with the Astral species tree, with strong support for nodes separating North American from Eurasian + African clades, while the vast majority of remaining nodes were poorly supported (Fig. 1d). The mean crown age of *Colias* was estimated at 2.66 million years old (2.10 – 3.34 posterior distribution 95 % limits), while the mean age of the last common ancestor of the non-South American *Colias* at 0.98 million years ago (0.75-1.21). The SNAPP analysis also estimated extremely short branch lengths among the majority of species (Fig. 1d), which corresponds to the extensive conflict among gene trees (Fig. 1a) and previous single and multigene studies (Limeri and Morehouse, 2016; Pollock et al., 1998; Wheat and Watt, 2008), in suggesting that non-South American *Colias* rapidly diversified into two regional clades in the past million years. Using these results, we can now view Alba and putative Alba phenotypes in a global phylogenetic context, revealing that Alba is in all the geographic clades of *Colias* (Fig. 1b, d). While much of the phylogenetic conflict observed arises from rapid speciation events and incomplete lineage sorting, introgression has likely been extensive during these events and potentially an important contributor to moving Alba among species and regions.

### Genome-wide Introgression analysis

In order to assess to what extent introgression played in the current distribution of Alba, we first evaluated introgression among *Colias* species using D-statistics. All possible species trios were assessed for introgression, using the South American *C. lesbia* as an outgroup. Grouping results by region (North America, Eurasian, Holoartic), we found significant levels of introgression among non-South American species (Fig. 1e, Fig. 2). Additionally, the proportion of significant trios showing introgression, as well as the general level of introgression, increased when we combined taxa from either North America or the Eurasian region with the Holoartic taxa. To estimate when these introgression events among the non-South American species occurred, we combined our D-statistics with our species tree and used the F-branch metric (Malinsky et al., 2018), which differentiates signatures of historical introgression between internal branch nodes from introgression between extant species. This revealed introgression between an ancestor of the Holarctic *nastes* clade and the North American species, as well as low levels of introgression among the Eurasian species (Fig. 2). Since species in the *nastes* clade are fixed for Alba, the Alba allele may have been transferred through introgression between this clade and an ancestor to the North American species. However, this analysis is unable to resolve the direction of introgression or capture localized intra-chromosomal introgression events. To specifically investigate the role of introgression around the Alba locus, sliding windows of 50 SNPs across the genome were analyzed and f_dM calculated (an alternative statistic to D that is appropriate for windows-based analyses). However, no significant signatures of elevated introgression near *BarH1* were detected in any species trio (Supplementary Fig. 3). While this suggests no introgression of the *Alba* locus among *Colias* species, our results may arise due to a lack of sufficient informative SNPs to detect localized introgression. In *C. crocea*, Alba is caused by transposable element insertion not found in orange haplotypes. In our f_dM analysis, this Alba insertion is necessarily excluded when analyzing all species. Thus, our intra-chromosomal analysis of introgression is dependent on finding enough linked variants in the region surrounding the Alba locus rather than the locus itself, and this could reduce analysis power. Thus, our failure to detect local introgression dynamics near the identified Alba locus in *C. crocea* could be due to insufficient power resulting from the genomic architecture of the trait, introgression not moving the Alba locus extensively or recently among *Colias* species, or Alba having a different genetic basis among species.

**Figure 2.**
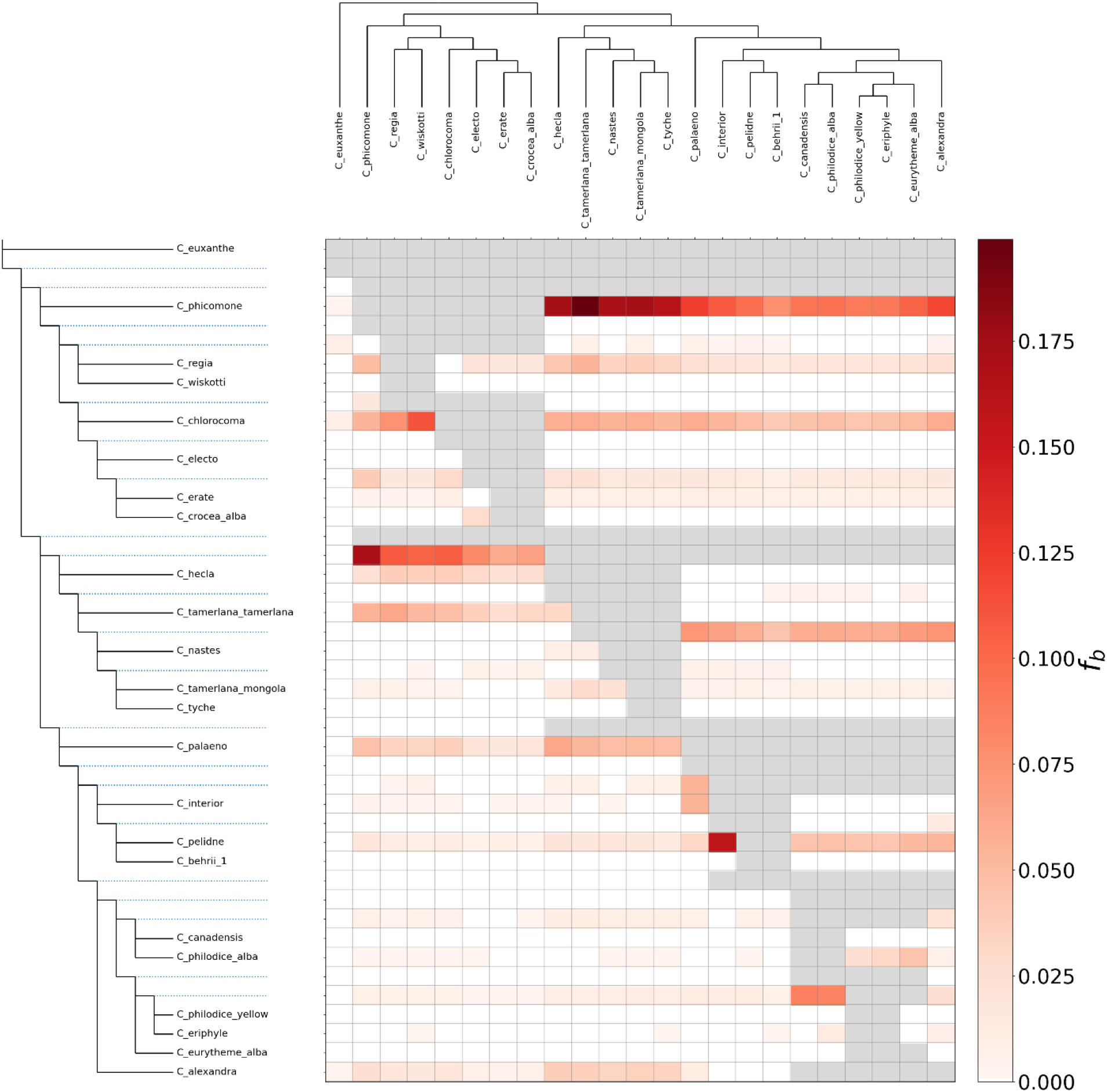
Signature of historical introgression across the *Colias* species tree phylogeny. Each cell in the grid indicates the F-branch statistic, identifying excess sharing of derived alleles between branch nodes on the y-axis and individual species on the x-axis. A darker color in the heatmap indicates higher F_B_ and suggests gene flow between that branch and species. Allele sharing between an internal node and current species can be used to infer past gene flow events between an ancestor of a clade A and a species B. Results indicate a strong signal of introgression between an ancestor of the *nastes* clade and the North American clade, as well as between *C. phicomone* and the Eurasian species.

### Identification of the Alba locus in Colias eurytheme

In order to thoroughly test whether Alba has a shared or *de novo* origin among species, we next mapped Alba in the North American species, *Colias eurytheme*. We first identified the chromosome carrying Alba, using a linkage map generated from a *Colias eurytheme* x *Colias philodice* F2 cross segregating the Alba phenotype, which allowed us to not only position our genome assembly scaffolds in chromosomal order but also to identify the chromosome bearing the Alba locus (Fig. 3a). This F2 cross showed Mendelian 1:1 segregation of the dominant Alba phenotypic state among the F2 females (*χ*^2^=0.006, d.f.=2, P-value>0.9), showing inheritance from an F1 single hybrid parent. The linkage map positioned the Alba locus on Chromosome 3 (Fig. 3b). Unfortunately, we could not further resolve the position of Alba within this chromosome since the Alba polymorphism was inherited from the maternal side of the F1 cross and was thus void of crossing-over events, due to the achiasmatic mode of maternal meiosis in Lepidoptera (Traut et al., 2007).

**Figure 3.**
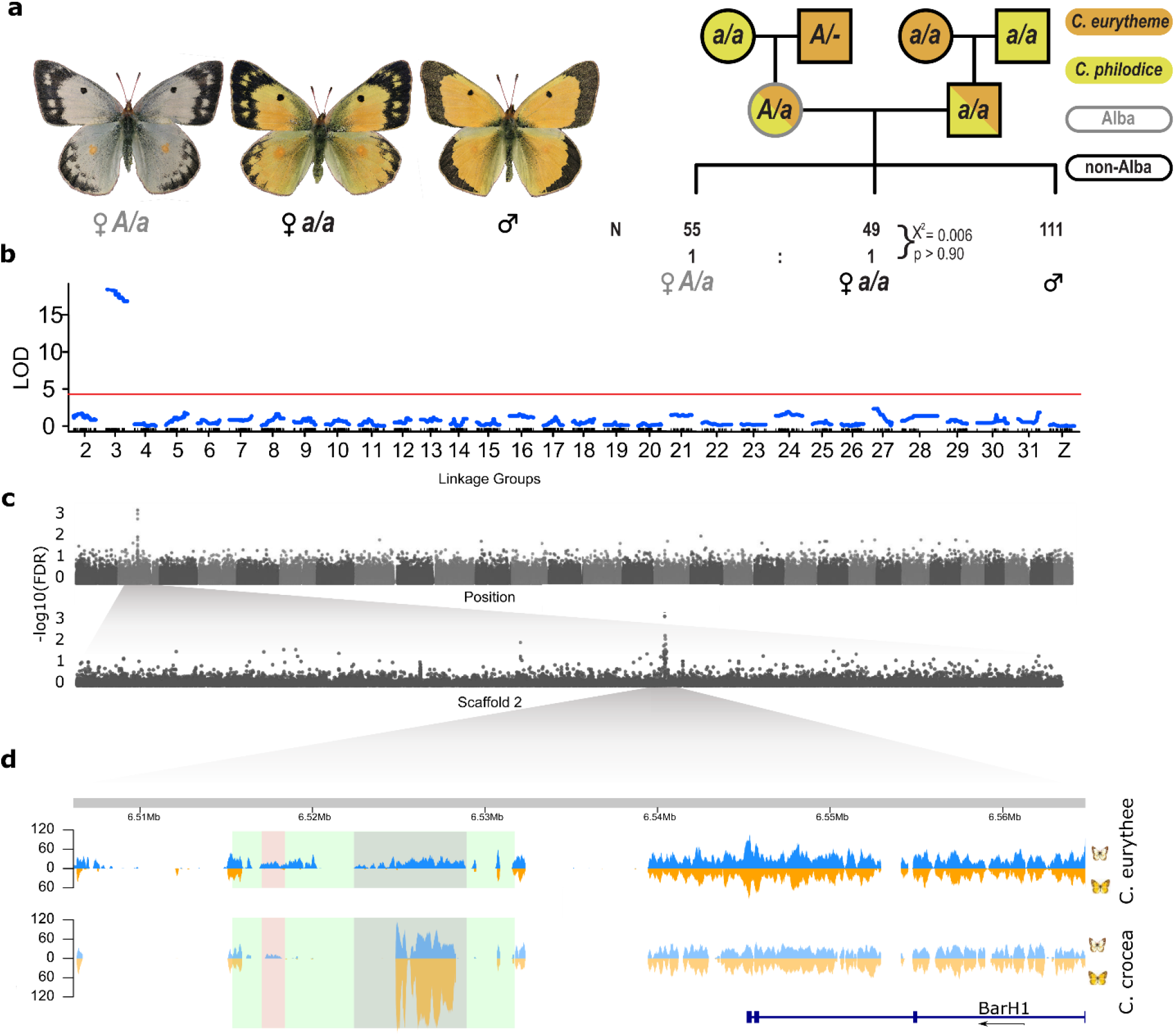
Identification of the *Alba* locus in *Colias eurytheme*. **a.** *Colias eurytheme* specimens and schematic of female informative crosses used in the linkage analysis. **b**. Linkage mapping of the Alba trait generated from female informative hybrid crosses of *C. eurytheme* and *C. philodice* revealed a single autosomal locus on Chr. 3 that was associated with Alba. **c.** GWAS using WGS data from 14 and 15 orange and Alba *C. eurytheme* females. Blocks in the manhattan-plot are colored by chromosome, ordered 1:31 (y-axis represent negative log value Bonferroni-Holm corrected false discovery rate (-log10(FDR_BH)) of each variant, the x-axis is scaffold position, the second row is a close up of scaffold 2, which makes up part of Chr. 3 as well as containing the *Alba* locus. **d.** Detailed view of the 58kb region identified from the 10X Chromium assembly showing WGS mapping coverage of an Alba (blue) and orange (orange) female from *C. eurytheme* (top) and *C. crocea* (bottom), respectively. The reads were aligned to the entire Alba reference genome (filtered to MAPQ > 20 and good pairs). Colored boxes highlight the *Alba* insertion (green), repetitive content within the insertion (grey), and a region of high sequence similarity with *C. crocea Alba* insertion, which is referred to as the *Alba* candidate locus (red). The grey region unique to Alba individuals in *C. eurytheme* is not unique to Alba individuals in *C. crocea* and is found at elevated coverage in all Eurasian species. Location and direction of the *BarH1* gene is indicated by the blue gene model.

A genome-wide association study (GWAS) was used to fine-map the Alba locus within *C. eurytheme*, using genomic data from 15 Alba and 14 orange wild-caught females mapped the reference genome. This identified two loci, the most significant of which was a single locus on Chr. 3 situated immediately downstream of the *BarH1* gene (Supplementary Fig. 4; but also Fig. 3c,d), which is concordant with both our previous mapping (Fig.3b) and the location of the Alba locus identified in *C. crocea* (Woronik et al., 2018). The second locus, which had less support, was located on a different chromosome between a PIFI-like helicase and a PiggyBac transposon (Supplementary Fig.5). We hypothesized that the second locus was an artifact arising from aligning reads to a reference genome lacking the Alba insertion since the reference was from an orange female individual. To test this, we generated an Alba genomic reference by combining a draft assembly made using linked read technology from an Alba *C. eurytheme* female and the reference genome (Supplementary Fig. 6), which added ~36kb of sequence downstream of *BarH1*. Repeating the GWAS using this synthetic Alba reference genome identified only the previous *BarH1*-associated locus, indicating reference bias as a likely cause of the second peak (Fig. 3c).

### Investigating the Alba insertion

To further investigate the Alba associated insertion region, which we expected to be composed of repeat content and regions unique to Alba, we conducted a read depth analysis by mapping the *C. eurytheme* individual genomes from the GWAS onto the Alba genome. By contrasting uniquely mapped reads that were at expected coverage depth to 1) reads mapping at higher-than-expected depth, or 2) reads not mapping uniquely, we could distinguish between unique Alba content and low complexity or repeat regions found in other parts of the genome. In the Alba associated insertion region, we identified an approximately 20 kb region containing two stretches of unique Alba content (where no orange reads mapped); data from orange females showed no unique content (Fig. 3d). Next, we similarly aligned reads from orange and Alba *C. crocea* females (n=15 each), revealing that only one of these two regions contained reads unique to Alba in both species (Fig. 3d), suggesting a region of high sequence similarity between the two species, which is notable given their deep divergence (Fig. 1d). We hypothesized that this shared region causes the Alba ALHS and hereafter refer to it as the Alba candidate locus (Fig. 3d). We further documented the uniqueness of the Alba candidate locus in *C. eurytheme* using PCR genotyping for an additional eight wild-caught females of each color morph (Supplementary Fig. 7).

### Comparative analysis of Alba candidate locus

To test the hypothesis that our identified Alba candidate locus is associated with Alba in additional *Colias* species, we generated an additional draft genome to resolve the Alba locus for *C. nastes*, a species fixed for the Alba color phenotype. In *C. nastes*, the Alba candidate locus and the *BarH1* gene assembled as a single contig, consistent with the hypothesis of the genetic basis for Alba having a shared ancestry among *Colias* species (Supplementary Fig. 8–9). In the aforementioned second region, which was unique to Alba only in *C. eurytheme*, we observed a dramatic increase in read depth and nucleotide diversity in *C. crocea*. This suggests that this segment is found in more than one copy in both orange and Alba *C. crocea* individuals and highlights the complex nature and evolutionary history of the locus.

Next, we formally tested the hypothesis of a shared origin of Alba by quantifying the association between having the Alba candidate locus and the Alba phenotype. All species with white wings had reads covering the entire Alba candidate locus except for *C. phicomone*, where reads covered only approximately half the candidate locus. In contrast, none of the samples from females with colored wings had reads covering the insertion region, instead reads piled up in low complexity areas flanking the locus. Thus, we observed a perfect correlation among species between having the insertion and the Alba phenotype. To estimate the likelihood of such a correlation on a genomic scale across species, we performed a window-based analysis of read coverage across the genome for all species (n=546,228 windows, 600bp in length each). A window located in the Alba candidate locus was the only region in the entire genome where the presence of coverage segregated with female wing color (Supplementary Fig. 10), suggesting the Alba insertion causes Alba across *Colias*.

### Phylogenetic analysis of the Alba locus

To further investigate the evolution of the Alba candidate locus, we constructed a phylogenetic tree for this region, which revealed a grouping of species discordant with the species tree (Fig. 5). While the *nastes* clade, European *C. crocea* and *C. erate* were all placed concordantly with the species tree, North American samples were not. Instead, they showed a pattern suggestive of separate introgression events. *C. eurytheme* grouped with *C. philodice* originating from the Maryland hybrid population and *C. nastes* clade as the closest outgroup, suggesting that Alba has been exchanged between the two former species during their ongoing hybridization and that this allele might have originated from an ancient introgression event with the Holarctic *C. nastes* clade. Meanwhile the *C. philodice* originating from British Columbia grouped with *C. canadensis and C. pelidne*, suggesting an independent introgression event of Alba entering *C. philodice* (though whether this sample is *C. philodice* or a subspecies thereof will require more regional sampling). Within *C. eurytheme*, our sampling of many Alba females reveals branch length differences among individual alleles, which suggest that the Alba allele has been maintained within *C. eurytheme* long enough for mutations to accumulate (Fig. 5). In contrast, the multiple alleles sampled from European *C. crocea* form a polytomy with *C. erate*, with the *C. erate* allele identical to several *C. crocea* samples, consistent with documented ongoing hybridization between *C. crocea* and *C. erate* (Descimon and Mallet, 2009). Together, these observations suggest that, while some regional hybridization has resulted in a reticulate phylogeny of the Alba locus, balancing selection has also played a role in maintaining Alba within species.

### Functional validation of insertion

Our comparative analyses of the Alba insertion locus strongly suggested that the conserved Alba locus has a functional role in regulating the expression of *BarH1* in females’ wing scales, so we tested this hypothesis by generating a somatic deletion mosaic of the Alba candidate locus using CRISPR/Cas9 in *C. crocea*. Along with Cas9, four gRNAs all targeting different parts of the locus were injected individually and together as a cocktail to generate multiple cuts and remove a significant portion within its 1.2 kb length. While injections with single gRNA did not produce any phenotypic changes (Supplementary Table. 7), of the forty eggs injected with the four-gRNA cocktail, unfortunately, due to bad rearing, many died before we were able to place the hatched larvae on hostplant, and we were only able to rear five individuals, two of which were males. From the remaining three females, we had two mosaic mutant individuals that exhibited extensive wing clones where scales recovered the orange pigmentation of non-Alba females (Fig. 4, Supplementary Fig. 11). Both females were genetically Alba, as validated by PCR (Supplementary Fig. 12, and the remaining orange female showed no abnormalities. Successful mutagenesis was confirmed by PCR fragment size polymorphism relative to uninjected Alba females. The observed wing phenotypes are reminiscent of the effects seen in previous mosaic knock-outs within the coding region of *BarH1* (Woronik et al., 2019). While the previous coding knock-outs produced eye color phenotypes in both sexes (Woronik et al., 2019), consistent with the known role of BarH1 in insect eye development (Hayashi et al., 1998; Kojima et al., 2000; Woronik et al., 2019), our work here targeting the Alba candidate locus did not produce any noticeable eye aberrations (Supplementary Table. 7), indicating it functions not only as a regulatory region but as a modular enhancer region required for female and scale-specific expression of *BarH1*.

**Figure 4.**
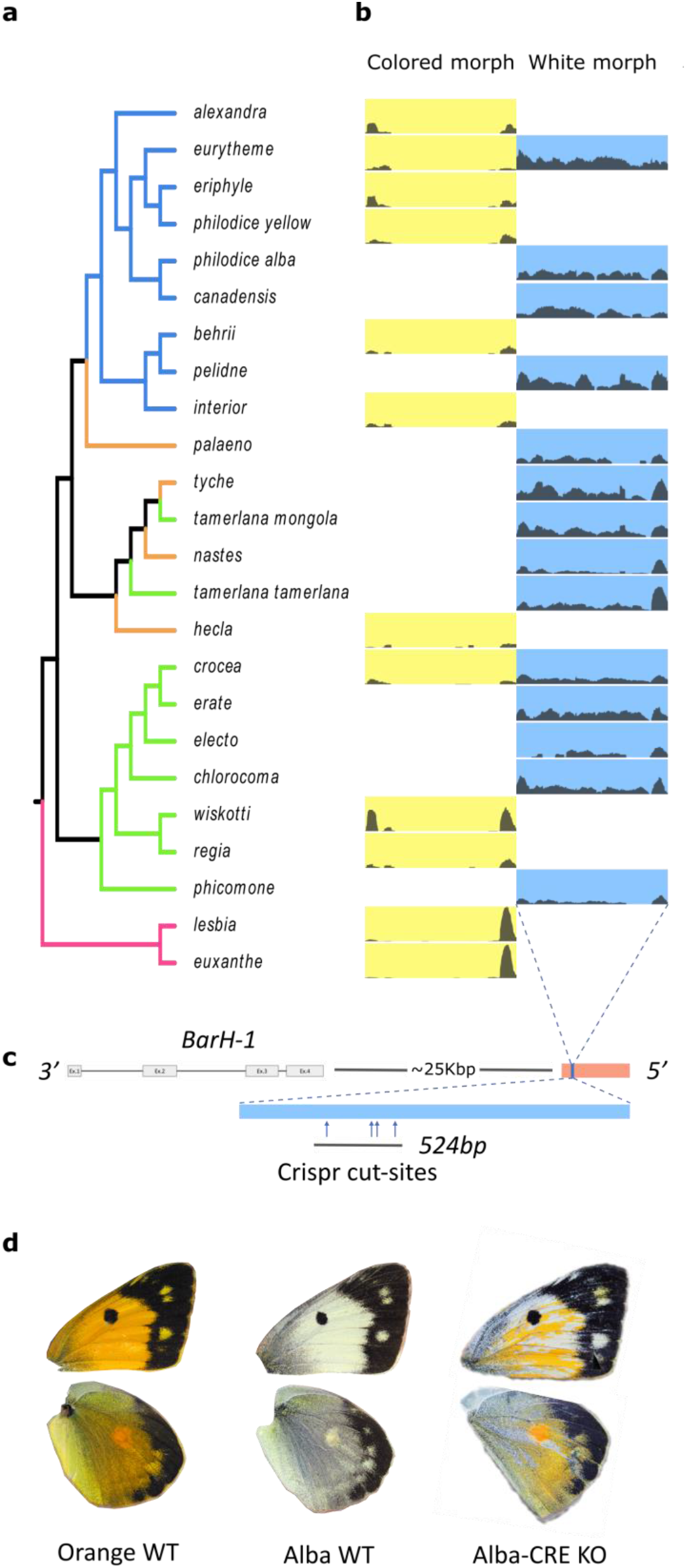
Association of the *Alba* candidate locus with wing color across a species tree of *Colias* butterflies. a. Species tree of *Colias* colored by geographic region, with purple = South America, blue = North America, orange = Holarctic, green = Eurasia and Northern Africa. b. Read depth coverage across the *Alba* candidate locus for the sequenced samples, with separate columns depicting coverage plots according to female wing color. In cases where we have sequence data for both morphs, both are shown next to each other. **c.** Location of the *Alba* candidate locus in *C. crocea* (blue square, in the *Alba* insertion (red) and location of CRISPR cut sites (blue arrows). **d**. Images of dorsal front- and hind-wings wild-type *C. crocea* females; from left to right: orange, Alba, and one of the successful *Alba-CRE* mutants. Orange areas on the wings of the on the *Alba-CRE* KO individual are somatic CRISPR mosaic KO cells showing a return of color production.

## Discussion

The distribution of the Alba ALHS within the *Colias* genus, wherein some species are polymorphic for Alba or fixed for either Alba or the color phenotype, has been facilitated by a complex interplay between balancing selection and introgression. By identifying and comparing the genetic basis of Alba in two species that last shared a common ancestor at the base of the main species radiation in *Colias, we* confirm Alba’s single ancestral origin for most, if not all *Colias*. Our comparative analysis of this trans-specific polymorphism allowed us to hypothesize about the location of Alba’s alternate ***cis***-regulatory module of *BarH1*, which we were able to confirm by genetic manipulation. The resulting indel-associated regulatory sequence, which has a wing-specific effect on the function of the transcription factor *BarH1*, is in concordance with the hypothesis that discrete regulatory modules with spatially discrete functions might be common in morphological evolution (Prud’homme et al., 2007). Our findings extend this hypothesis to an ALHS and constitute the first functional identification of a modular enhancer in a butterfly.

For the majority of *Colias* species, we are able to reject the hypothesis of multiple independent origins of Alba. While we cannot identify the maximum age of Alba at this time, we show it evolved prior to the separation of the North American and Eurasian clades approx. one million years ago. Additionally, as most *Colias* have at least one generation per year, this polymorphism has been maintained in these divergent clades since they shared a common ancestor at least one million generations ago. However, given the presence of Alba in South American taxa (Hovanitz, 1945), Alba is likely to be much older (e.g., age of *Colias*). Thus, the Alba ALHS is an ancient trans-specific polymorphism. The larger challenge is determining the relative role ancient balancing selection and introgression have played in its maintenance over time (Gao et al., 2015). Within-species diversity of the *Alba* allele is consistent with balancing selection (Fig. 5). While we do find evidence of historical introgression events (Fig 1c, Fig. 2), we did not detect any significant localized introgression around the *Alba* locus (Supplementary Fig. 3), potentially due to low power arising from the genetic architecture of *Alba*. However, the *Alba* locus phylogeny (Fig. 5) suggests several recent introgression events that include the *Alba* allele (Fig. 1).

**Figure 5.**
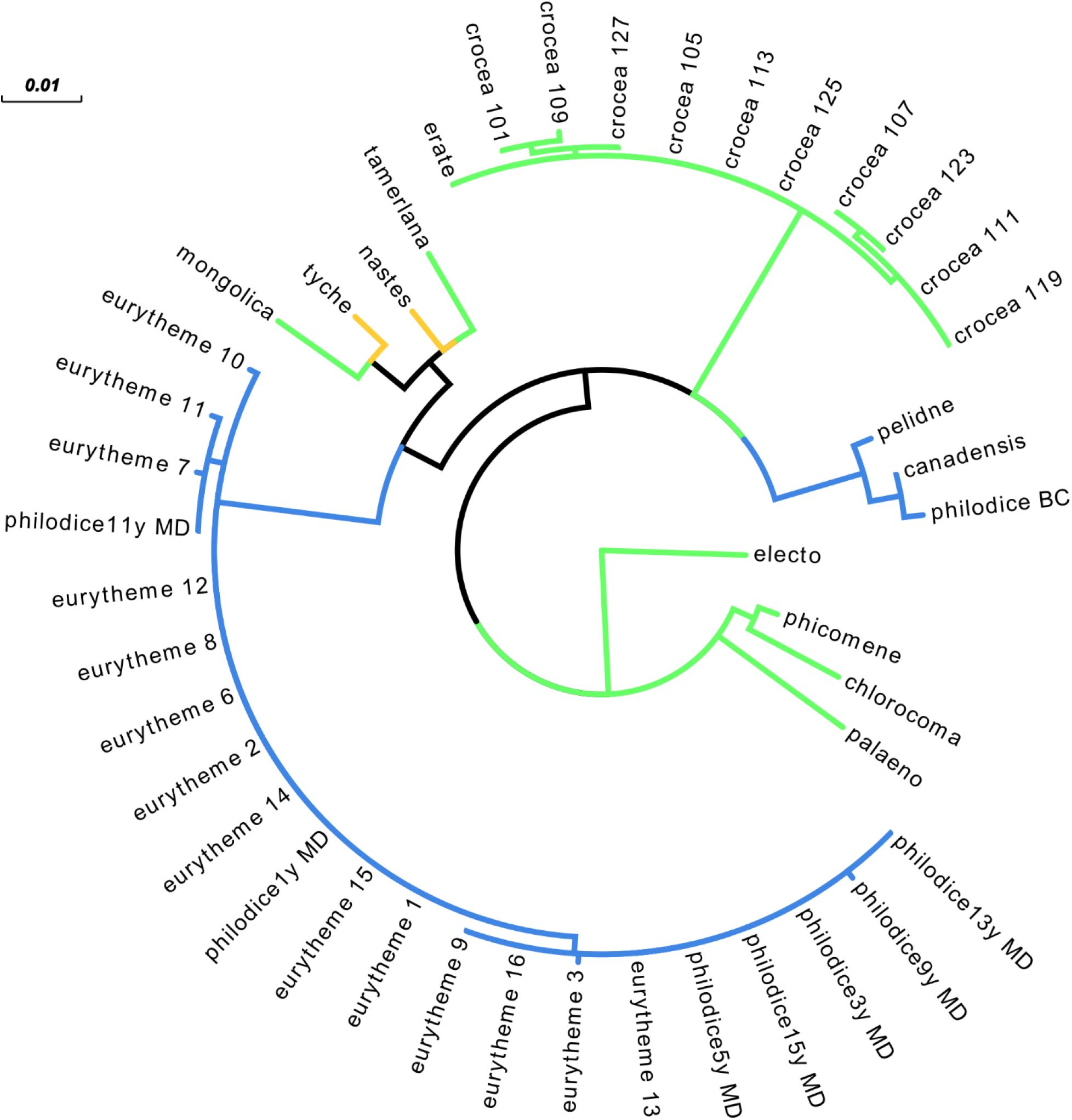
Phylogeny of the 1.2 Kbp long *Alba* candidate locus. Branch color corresponds to geographic region (blue = North America, yellow = Holarctic, green = Eurasia and Northern Africa), and branch length corresponds to substitution differences. The branching structure reveals shared alleles among *C. eurytheme* and the *C. philodice* originating from the Maryland hybrid population. Note the distinct variation among alleles in *C. eurytheme*, which suggests these alleles have been within this species long enough to accumulate variation. While similar diversity is seen within *C. crocea*, the shared allele with *C. erate* suggests ongoing Alba allelic exchange between these species, similar to that between *C. eurytheme* and *C. philodice*. The large divergence between the alleles found in the Maryland *C. philodice* and the *C. philodice* used in the phylogenetic analysis further supports the reticulate evolution of Alba among North American *Colias*.

The study of the genus-wide maintenance of an ALHS presented us with many inherent challenges that may account for the paucity of other well-characterized examples in the literature. Some ALHS may be exclusively physiological and lack a clear visual manifestation, which makes tracking the presence or absence of the ALHS across taxa particularly challenging. Our study here relied upon the large body of work linking resource allocation and performance with wing color (Gilchrist and Rutowski, 1986; Graham et al., 1980; Limeri and Morehouse, 2016; Nielsen and Watt, 1998; Remington, 1954; Watt, 1973; Woronik et al., 2018, 2019), allowing us to include additional species where the specific metabolic resource allocation tradeoffs of Alba have yet to be established. Furthermore, to persist through time, polymorphisms of complex traits, such as ALHS, are expected be the result of structural variants, as these are less likely to be broken up by recombination (Llaurens et al., 2017). However, the genetic tools traditionally used for the identification of genetic associations generally exclude, and lack power to accurately detect, structural variants (Chaisson et al., 2019; Kishikawa et al., 2019; Mérot et al., 2020b). If the causal locus is absent in the genomic reference, as was the case when we used our orange reference genome for our GWAS analysis, causally associated reads cannot map to the genome, while repeat content within the causal locus will map to non-orthologous regions producing spurious associations (Supplementary Fig. 4, 5). Additionally, since downstream analyses and phylogenetic tools generally require discarding indel variants (Alonge et al., 2020), this likely decreased our power to detect localized introgression near the *Alba* locus. Fortunately, individual resequencing using long reads is becoming more accessible and has already been essential for the discovery of novel genomic regions associated with adaptation, such as premating isolation and speciation in crows (Weissensteiner et al., 2020). Further development of tools and pipelines for the downstream analysis of structural variants via long read data at the individual level will hopefully remove the aforementioned biases.

Finally, while the use of CRISPR/Cas9 for the functional validation of candidate genes is now accessible to a wide range of taxa (Sun et al., 2017), when candidate gene KOs have lethal effects the generation of transgenic lines are impossible. Hence, many such studies have been limited to assessing effects in low penetrance F0 mosaic phenotypes (e.g. Woronik et al., 2019; Zhang and Reed, 2017). While such mosaic phenotypes have greatly advanced the study of morphological traits (Burg et al., 2020; Perry et al., 2016; Westerman et al., 2018b), such mosaics of physiological phenotypes are nearly impossible to interpret, leaving the study of non-morphological phenotypes unable to use these new advances. While targeting the cis-regulatory regions of genes isn’t a novel idea, the inherent difficulty in identifying such candidate regions have limited this approach. Here, the conserved nature of the *Alba* allele allowed us to leverage a phylogenetic foot printing approach (Tomoyasu and Halfon, 2020) to localize the locus and conduct the first targeted mutagenesis of a regulatory enhancer in Lepidoptera (but see (Murugesan et al., 2021)). Finally, while induced mutations in the coding exons of *BarH1* had a high lethality and pleiotropic effects (*e.g*. eye phenotypes), mutations in the CRE region of *Alba* were not lethal and appear to lack such negative pleiotropic effects. This is consistent with the CRE being a modular enhancer, only necessary for regulating *BarH1* expression in scale cell precursors. Facilitated by its modularity, introgression of this segment alone could be sufficient to (re)introduce *Alba* in a species lacking the Alba ALHS. Future studies in the system will be able to take advantage of these advances, especially those allowing the establishment of germline mutations, which will enable further detailed studies not only of the *Alba* CRE region, but direct investigation of the physiological aspects of the Alba ALHS.

## Methods

### C. eurytheme genome

High molecular weight DNA from six female pupae originating from Davis (CA, USA) and reared in the lab for several generations was sequenced on PacBio Sequel v1 at the U. Maryland - Baltimore Institute of Genomic Sciences. Assembly followed the Falcon/Falcon-Unzip/Arrow assembly pipeline (Chin et al., 2016) and led to a diploid genome length of 583Mb, with N50 of 2.7Mb, haploidization was performed using Haplomerger2 leading to a haploid genome of 364.5Mb and 123 scaffolds (Huang et al., 2017).

### C. eurytheme genome polishing, quality control, and annotation

Pilon v.1.22 (Walker et al., 2014) was used to polish the genome, using 150bp PE reads (350bp insert, Illumina HiseqX) aligned with NextGenMap v.0.5.2 (Sedlazeck et al., 2013). Genome quality before and after polishing was assessed using Busco v1.1b1 with OrthoDBs Lepidoptera v10, as well as N50 (Seppey et al., 2019; Simão et al., 2015). Repetitive regions were softmasked using RED v:05/22/2015 (Girgis, 2015). The genome was annotated using the Braker2 (Brůna et al., 2020) pipeline with transcriptome data generated in a previous study (Nallu et al., 2018) aligned with Hisat2 v2.2.1 (Kim et al., 2019) and protein data from OrthoDBs Arthopod database (V10).

### Synteny comparative analysis

To assess our genome assembly and check for any large-scale structural changes compared to other sequenced Lepidopteran genomes, we compared our *C. eurytheme* chromosome to one from the sister genus, *Zerene cesonia* (Rodriguez-Caro et al., 2020). Whole-genome alignments were performed using nucmer v4.0 (Marçais et al., 2018) followed by circos plotting using the R package circlize v.0.4.9 (Gu et al., 2014).

### Colias phylogenetic analysis

For each individual, whole-genome sequencing reads were generated via DNA extracted from thorax and/or abdomen via a salting out method (Aljanabi and Martinez, 1997). DNA quality was evaluated using a 260/280 ratio (Nanodrop 8000 spectrophotometer; Thermo Scientific, Waltham, MA, USA). The library preparation and short read paired end sequencing (500Bp insert) for all individuals was performed at BGI China. Reads were filtered for adapters and trimmed at the 5’ and 3’ end based on a PHRED quality score >20. Reads were aligned to the *C. eurytheme* reference genome using NextGenMap v0.5.5 (Sedlazeck et al., 2013). Using these bam files after MapQ > 20 filtering via Samtools v.1.9 (Li et al., 2009), the longest exon per gene for each individual was obtained the CDS annotation for *C. eurytheme* via bam2fasta script from the package bambam v1.4 tool-kit (Supplementary methods)(Page et al., 2014). *Zerene cesonia* (Rodriguez-Caro et al., 2020) was used as an outgroup, the dataset of which was generated by aligning Illumina sequencing reads from *Z. cesonia* (SRR11021459) to the *C. eurytheme* genome, as per the bam2fasta pipeline outlined above. Individual gene trees were then estimated using iQTree v. 2.0.6, which were then used to estimate a species tree via ASTRAL v.5.7.3 (Zhang et al., 2018). Gene tree support for the species tree was assessed using Phyparts (https://bitbucket.org/blackrim/phyparts/src/master/). Species trees for each chromosome were generated using all genes trees of a given chromosomes to generate an Astral species tree. Analyses were repeated after higher quality filtering, with identical species tree results (Supplemental methods). SNAPP v.1.3.0 (Bryant et al., 2012) analysis, implemented in BEAST2 v. 2.6.3 (Bouckaert et al., 2019), followed previous extensive analyses for optimal analysis settings (Stange et al., 2018), with dataset construction using snapp_prep.rb (https://github.com/mmatschiner/snapp_prep). Calibration for the timing of the split between *Zerene* and *Colias* used a secondary calibration of 10.9 million years ago (Chazot et al., 2019), along with two monophyletic constraints set to increase run speed (SA taxa, non-SA taxa). Please see Supplementary methods for more details.

### Introgression analysis

Introgression between different species was estimated using D-statistics calculated from ABBA-BABA between all possible trio combinations using the Dsuite software package (v0.3)(Malinsky et al., 2020). Using Dsuite we also calculated an f-branch metric, a statistic related to the f-4 statistics, which allows you to summarize the amount of shared introgressed material on a branch and infer past gene-flow (Malinsky et al., 2018). Using the Alba reference genome, we aligned the reads of each species using NextGenMapper, and then called variants using Freebayes (Garrison and Marth, 2012). The resulting vcf-file was filtered (see SM for details) using vcftools (Danecek et al., 2011).

Using the Dinvestigate tool part of the Dsuite tool-kit f_d and f_dM was calculated in windows non-overlapping windows of 15 and 100 informative SNPs to look for signals of adaptive introgression along the chromosomes. (more details in Supplementary Methods)(Martin et al., 2015).

### C. eurytheme x C. philodice 2b-RADseq genotyping and linkage map

By generating a linkage map, using the 2b-RADseq (Wang et al., 2012)whole-genome genotypes of the F2 brood from a *C. eurytheme* x *C. philodice* hybrid cross, we turned the haploid genome scaffolds into chromosome-wide super-scaffolds. Linkage mapping followed the basic LepMap3 with some exceptions (explained in the SM) and resulted in 31 linkage groups, with one short unplaced scaffold. The linkage map was output as a 4-way cross, which was imported into R package r/qtl using custom code (contributed by Karl Broman). We performed a genome scan with a single QTL binary model and ran a permutation test (n=1000) to determine a 5% significance threshold.

### GWAS of Alba in C. eurytheme

Individuals used in the genome resequencing were from 15 Alba and 14 orange *C. eurytheme* females caught in Tracy, California, stored at −20 in 95% ethanol. For DNA extraction through to read cleaning, see Supplemental Materials. Cleaned reads were mapped to the *C. eurytheme* reference genome using NextGenMap v0.5.2, followed by duplicate marking, and then Freebayes v1.3.1-16-g85d7bfc for variant calling. The variants were filtered using VCFTOOLS v0.1.13 (Danecek et al., 2011). Variants were associated with the Alba phenotype using PLINK v1.9 (Chang et al., 2015). Two separate sets of filters were used, one with stronger priors, where the nature of the inheritance pattern was taken into account, and one weaker where sites were filtered quality and depth mainly; for more detailed information on the filters, please refer to the SM.

### Generation of an Alba specific reference genomes

To get an idea of what the sequence and structure of the Alba insertion was, we generated an Alba reference genome. First, we generated a draft genome using a 10X Chromium library, sequenced on a NovaSeq S4, 2×150bp PE reads, followed by assembly with Supernova v2.1.1 (performed by SciLifeLab). In addition to *C. eurytheme*, we also generated draft genomes for *C. nastes*, a species fixed for Alba as well as *C. crocea* (used as a control to compare against the previous genome), using the same protocol.

### Characterizing the Alba insertion in C. eurytheme

We identified the scaffold containing *BarH1* by using tBLASTn in the *C. eurytheme* Alba Supernova-assembly. We then aligned all the resequencing data from the GWAS to this contig. Read depth along the contig was analyzed visually in IGV, and differences between the Orange and Alba morph noted. Regions where no orange reads aligned, but Alba did, were extracted and blasted back against the *C. crocea* reference genome (Woronik 2019), to assess whether this was the previously identified Alba insertion region. To identify borders of the Alba insertion in *C. eurytheme*, we aligned the Alba contig against the orange *C. eurytheme* reference genome using BLASTn. This then provided the boundaries of the Alba insertion region for *C. eurytheme*, which was then used to place this haplotype into the orange *C. eurytheme* reference genome assembly, creating what we refer to as the *C. eurytheme* Alba reference genome.

### GWAS using the Alba reference genome

Using the new Alba reference genome, we repeated the steps done in the initial GWAS, seeing if the alternative loci disappeared with new targets to map against.

### PCR-based validation of insertion

The presence and uniqueness of the insertion to Alba individuals were validated using PCR-based markers with primers designed to bind within the insertion region. Primers were designed using primer3 software (libprimer3 release 2.5.0, Untergasser et al. - 2012). DNA from 8 orange and 8 Alba females independent females were used in the analysis, and CytC was used as a positive control in each reaction. For more information about primer design and the reaction, see the SM.

### Identification of Alba insertion in C. nastes

The genome assembly were scanned for the presence of the *C. crocea* insertion sequence, as well as the sequence identified in the *C. eurytheme* Chromium assembly, and whether it was found in linkage with the *BarH1* gene using the same combination of tBLASTn and BLASTn as we used in *C. eurytheme*.

### Alignment and assessment of the Alba insertion across species

Resequence-data generated for the phylogeny was aligned to the Alba reference using NextGenMap, filtered (MAPQ 20 and proper pairs), and had coverage across the Alba insertion region visually inspected in IGV (v2.7). We assessed whether the presence of read-coverage segregated with female wing color. Regions that were unique to only white-colored species (putative Alba ALHS species) were considered to be conserved regions of the Alba insertion and likely causal for the phenotype. The likelihood of coverage segregating between the two color-morphs to this degree was estimated using a window-based analysis (See SM).

### Phylogenetic relationship of Alba insertion

We extracted the consensus sequence of this using the bam2fasta tool (part of the bambamv1.4 *tool-kit*). We then assessed their phylogenetic relationships using IQtree with the same settings as in the primary analysis (Supplemental Methods).

### CRISPR/Cas9 targeted mutagenesis of the Alba insertion

We used the PROMO tool (Farré et al., 2003; Messeguer et al., 2002) to scan the conserved Alba locus for potential transcription factors, comparing the motifs against version 8.3 of the TRANSFAC database. The output of this scan was analyzed in three ways, 1) relevant candidate transcription factors were identified, such as *Doublesex;* 2) sites with a high density of potential TF-binding sites were recorded; and 3) sites that were highly conserved between different species were preferentially selected. We designed four sgRNAs seeking to produce multiple cuts and produce a large >100 bp deletion. gRNA design and injection were performed following the steps outlined in Woronik et al. 2019 and injected as a cocktail together with Cas9 at a 500 ng/ul concentration. *C. crocea* Alba females (n = 6) from Aiguamolls de l’Empordà, Spain, were captured and transported to Stockholm alive and allowed to oviposit on *Vicia villosa*. Eggs were collected three times daily for injection, ensuring that they were not more than 4 h old at the time of injection. Injected eggs were kept on glass slides inside sealed Petri dishes, together with moist paper. Hatched larvae were transferred to fresh *Vicia villosa* and kept in feeding cups with no more than five larvae at 23°C until pupation. Once pupated, the pupae were transferred to a climate cabinet kept at 16°C until eclosion. Mutated individuals had the CRISPR cut site validated using PCR-based assay.

## Supplemental Information

### Supplementary Methods

#### Bioinformatic Scripts

Detailed scripts for bioinformatic analysis, if not specified in the relevant section, be found on the projects github.

#### Colias eurytheme DNA extraction and genome sequencing

A *Colias eurytheme* stock originating from Davis (CA, USA) was maintained in a laboratory setting for several generations. High molecular weight DNA from six female pupae was isolated using the Qiagen Genomic-tip 100/G. The specimen yielding the most DNA (574.4 nanograms per microliter) was submitted for quality control, BluePippin extraction fragments > 15kb, PacBio SMRTBell Express library preparation, and PacBio Sequel v1 sequencing at the U. Maryland – Baltimore Institute of Genomic Sciences. Six PacBio Sequel v1 cells were sequenced and yielded a total of 50.46 Gb of sequence data with an average sub-read length of 10kb, *i.e*., a coverage of 144x assuming a genome size of 350 Mb based on fluocytometry. Assembly of the data using the Falcon/Falcon-Unzip/Arrow assembly pipeline (Chin et al. 2016) was outsourced to DNAnexus (Mountainview, CA), which led to a diploid genome length of 583 Mb with an N50 of 2.7 Mb. Haploidization of the genome was performed using Haplomerger2, leading to a haplogenome of 364.4 Mb assembled into 123 scaffolds with a scaffold N50 = 4.82 Mb.

#### C. eurytheme x C. philodice 2b-RADseq genotyping and linkage map

To further improve the assembly into chromosome-wide super-scaffolds, a linkage map of the genome was generated using the 2b-RADseq ((S. Wang et al. 2012)) whole-genome genotypes of the F2 brood from a C. *eurytheme* x *C. philodice* hybrid cross (B. Wang and Porter 2004). Briefly, we extracted DNA from the thorax of frozen individuals using a bead-shaker and the Quick-DNA 96 Kit (Zymo Research), digested 400 ng of DNA per individual with the *BcgI* enzyme (New England Biolabs), purified the 36 bp restriction fragments using the <u>ZR-96 Oligo Clean & Concentrator</u> kit (Zymo Research), added barcoded adapters corresponding to a 16 reduced tag representation (S. Wang et al. 2012) before ligation with the NEBNext Multiplex Primers Set 1-4 (New England Biolabs). The pooled library was enriched for a 155-175 bp (target inserts of 166bp) using a BluePippin instrument and sequenced using an Illumina HiSeq4000 SR50 run.

We investigated whether the haplogenome still contained some haplotype scaffolds/contigs. To achieve this, the two first steps (hm.batchA1.initiation_and_all_lastz + hm.batchA2.chainNet_and_netToMaf) of Haplomerger2 (Huang, Kang, and Xu 2017) were run on the haplogenome to create the alignment chain (all.chain.gz). The alignments of the remaining 123 scaffolds revealed that there were clearly some haplotypes left. We manually classified all scaffolds into full, partial, or unique scaffolds; there were 16 full and 11 partial haplotypes. Using a custom script, the remaining haplotypes and partial haplotypes were removed from the assembly to construct the final scaffold assembly with 108 scaffolds (one scaffold was cut to two, as likely chimera).

The genotype data for the linkage map for the *C. eurytheme* genome was obtained by Lep-MAP3 (LM3)(Rastas 2017) pipeline. First, the individual fastq files were mapped to the scaffold assembly using bwa mem (H. Li 2013), and using LM3 pipeline (pileupParser.awk, pileup2posterior.awk) and SAMtools mpileup (H. Li et al. 2009), we obtained the input genotype likelihoods.

Linkage mapping followed the basic LM3 pipeline as follows (non-default parameters inside parenthesis):

1. ParentCall2(ZLimit=2, removeNonInformative=1)
2. Filtering2 (dataTolerance=0.0001)
3. SeparateChromosomes2(lodLimit=14.5 maleTheta=0.5 femaleTheta=0.0001 distortionLod=1 sizeLimit=4)
4. 2 x JoinSingles2All (lodLimit=10 lodDifference=2 maleTheta=0.05 femaleTheta=0.0001 distortionLod=1)
5. OrderMarkers2 (chromosome=1..31 recombination2=0 informativeMask=13 useMorgan=1).

This yielded 31 linkage groups but after inspecting the markers occurring in the same scaffolds, two linkage groups were joined (27+30 and 29+25). Moreover, two linkage groups had very long maps (>100cM) and these groups were split, group 1 with SeparateChromosomes2 (map=map14.5.txt maleTheta=0.5 femaleTheta=0.0001 distortionLod=1 lg=1 renameLGs=0 lodLimit=16) and group 10 based on sex/autosome markers (indicated by * in the output of ParentCall2).

After these splits and joins, the JoinSingles2All and OrderMarkers2 were run again to obtain final (*de novo*) linkage maps with 31 linkage groups. The splits and joins were necessary due to complex family structure (multiple families) and low marker density.

With the help of the linkage map, the scaffolds (all except Sc0000116) were manually put together into these 31 linkage groups. Linkage groups were named based on chromosome numbers in *Melitea cinxia* (Ahola et al. 2014). The linkage map was re-evaluated in this physical order with OrderMarkers2 (recombination2=0 chromosome=1..31 evaluateOrder=phys_order improveOrder=0 hyperPhaser=1 phasingIterations=3), put into grand parental phase (phasematch.awk) and this map was used for QTL mapping.

#### C. eurytheme genome polishing, quality control, and annotation

Polishing of the genome was performed using Pilon v1.2.2 (Walker et al. - 2014), using data from a single orange female, with a Illumina TruSeq Nano library prep and sequenced (150 bp PE reads with 350bp350 bp insert size, Illumina HiSeqX) to provide ~30X genome coverage, aligned using NextGenMap v0.5.2. The assembly quality was assessed using custom scripts for basic length metrics and BUSCO v1.1b1 before and after polishing to evaluate the difference, using the insecta_odb9 dataset. The genome was softmasked for repetitive regions using RED (Version: 05/22/2015, Girgis, H.Z 2015 (Girgis 2015)).

Genome annotation was generated using BRAKER2 (v2.1) trained on data from C. eurytheme transcriptome and proteins. RNA-seq data generated in a previous study (Nallu et al.?), consisting or transcriptome data from several developmental life stages. We used a reference protein dataset from the arthropoda section of OrthoDB (v10). Transcriptome reads were aligned using HISAT2 v.2.1.0 (Kim et al. 2019, 2), against the unmasked genome, and the alignment was then filtered, sorted and indexed using SAMTOOLS v.1.7. Braker2 was run using the ETP mode and set to take softmasking into account. Quality of the annotation was then assessed by counting the number of good transcripts (genes containing both start and stop codon). Further assessment was done using OHR analysis (modified from (O’Neil et al. 2010))), reads transcripts with 0.9 identity or higher were clustered using CDHit v.4.8.1 (W. Li and Godzik 2006) and then compared against a protein set from B. mori. This gives an estimate of the fraction of full-length transcripts in the annotation, as well as completeness. Annotation of the resulting chromosome level assembly identified 18,460 genes, 18,081 of which had a correct start and stop codon. Clustering these “good genes’’ at 90% identity resulted in 16,352 genes. When we compared our annotation with the *B. mori* proteome, our set covered 11,623 of the 14,052 *B. mori* proteins with at least 80% of the *B. mori* length, indicating a high-quality annotation.

#### Synteny comparative analysis

To assess our genome assembly and check for any large-scale structural changes compared to other sequenced Lepidopteran genomes, we compared our C. eurytheme chromonome to one from the sister genus, Zerene cesonia (Rodriguez-Caro et al. 2020). Whole-genome alignments were performed using nucmer followed by circos plotting using the R package circlize v.0.4.9 (https://academic.oup.com/bioinformatics/article/30/19/2811/2422259). Bioinformatic details can be found on our github page.

#### Phylogenetic analyses

The longest exon per gene dataset was generated by running BUSCO upon the protein dataset from our annotation of the *C. eurytheme* genome, with the protein dataset generated using our GFF annotation and the genome as inputs for the gffread script from cufflinks v.2.2.1 (Trapnell et al. 2010). Of the total lepidopteran BUSCO genes searched (n= 5286), 4476 were found complete and single copy in our protein dataset for *C. eurytheme*. Among the BUSCO outputs is a table where for each annotated protein identified, its BUSCO status is indicated (e.g. as single and complete, duplicated, etc). Using this table, the exons of the complete and single copy proteins in our annotation were extracted and converted to bed. Then the length of each exon was calculated, allowing for the longest exon per protein ID to be selected using custom scripts, and the resulting bed file of these longest exns was then used as input for the bam2fasta script from the package bambam v1.4 tool-kit (Page et al. 2014), along with all the bam alignment files and a minimum depth requirement of 5 reads per base pair. The resulting set of fasta files (busco_exons) was then used to generate gene trees using iQtree, with each gene tree using extended model selection, a random starting tree, 1000 ultrafast bootstraps and optimization, and *Z. cesonia* set as an outgroup (-m MFP -t RANDOM -bb 1000 -alrt 1000 -bnni -o Z_cesonia). A total of 4244 gene trees were generated. These were then used as input for species tree estimate by Astral using default settings. Using the genome annotation, gene trees were also grouped by chromosome, which allowed for species tree for each chromosome to be similarly estimated.

We also generated a filtered set of the busco_exon dataset, using AMAS V.1.0 (Borowiec 2016) to remove *Z. cesonia* from all fasta files, and then generate a summary table of all files, which was then parsed to produce a set of files filtered to remove those with missing content > 1 %, < 5 % variable sites), and length < 300 bp and > 2000 bp. With this filtered set of fasts file IDs (n=1400), these iQtree gene trees were then selected for species tree estimation, to assess whether dataset quality had any effect on species tree topology. The resulting species tree from these filtered fasta files was identical to the full busco_exon analysis. Additional analyses using full CDS for ~9000 genes, or their 2^nd^ exon, produced essentially identical species tree results (data not shown).

Gene tree concordance with the species tree was assessed using Phyplots (Smith et al. 2015), to calculate the number of gene trees concordant and discordant with the species tree topology, per node. Phyplots output was further parsed to distinguish among gene trees discordant with the species tree, into those supporting a main alternative tree vs. many alternative trees, as well as those gene trees having less than 50% bootstrap support at the node in question. We used pieplots to represent these proportions (https://github.com/mossmatters/MJPythonNotebooks/blob/master/PhyParts_PieCharts.ipynb). Importantly, these pieplots results were concordant with estimated gene concordance factors via iQtree (data not shown).

SNAPP analyses (Bryant et al. 2012), which are run as an addon to the BEAST2 software program, were used for generating multi-species-coalescent analyses, but instead of using whole genes as in StarBEAST2, SNAPP uses SNPs. A further difference to StarBEAST2 is that gene trees are not estimated for each SNP, although SNPs are considered unique markers SNAPP; SNAPP estimate the species tree probability by integrating over all the possible gene trees observed among the SNPs. Despite this approach dramatically decreasing parameter space for analyses, this approach remains computationally demanding. Recent work has investigated the number of SNPs for optimal inference while minimizing computation demands, as well determined accurate SNAPP settings for time calibration, which uses a strick-clock model using fossil calibrations and the linking of all population sizes during analysis (Stange et al. 2018). Thus, in order to stay within a multi-species-coalescent framework for divergence time and phylogenetic relationship estimation, we followed these recommendations(Stange et al. 2018) and down sampled our taxa to remove redundancy among closely related species while retaining regional diversity. This took our full dataset of 29 species down to 18. Using AMAS, exon fasta files were subsamled to these 18 species, exons concatenated and then converted to phylip file format. A ruby script was used to generate an input XML file for SNAPP via snapp_prep.rb (https://github.com/mmatschiner/snapp_prep), which takes a phylip formatted sequence dataset, allows for specifications of run iterations, inter-SNP intervals, total SNPs, a starting tree, and diverse constraints to incorporated. Constraints included a temporal calibration for the timing of the split between *Zerene* and *Colias* using a secondary calibration of 10.9 million years ago with sigma=1, along with two monophyletic constraints well supported by previous Astral analyses (one for South American taxa, one for the remaining *Colias* species). Two SNP datasets were constructed, each containing 1000 random SNPs: 1) SNPs sampled from among the concatenated BUSCO exons with at least 300 bp between each SNP, 2) SNPs from among concatenated filtered BUSCO exons, with at least 100 bp between each SNP. Each dataset was run twice, for between 2 to 3 million iterations, which returned effective sample sizes (ESS) > 200 and were convergent within dataset with stable likelihood values, which was assessed using Tracer in the BEAST2 package. Posterior estimates used mean tree credibility after a 10% of data was discarded as burn n, as implemented in Treeannotator in the BEAST2 package. The resulting phylogenetic relationships and divergence estimates for *Colias* and non-South American *Colias* crown groups we nearly identical in their results, which was visualized using Figtree v.1.4.4 (Rambaut and Drummond 2012).

#### Introgression analysis

In order to assess the level of gene flow and introgression among Colias species, we calculated the D-statistic between all possible trios of species for which we had sequence data. Using the software Dsuite (ref) we were able to analyze this jointly and infer between which, past and present taxa, we’ve likely had introgression (Malinsky, Matschiner, and Svardal 2020). By calculating a F-branch statistic and analyzing it together with the phylogeny, we can infer between which taxa and which nodes in the phylogeny gene flow is most likely.

For this analysis, we used the sequence data for different *Colias* species mentioned previously and aligned them to the manually curated Alba-reference genome using NGM (same settings as previously). No filtering was done before using Freebayes to call variants. The resulting vcf-file filtered for strand-bias, quality score of 30, 90% of species sharing the site, minimum sample depth of 5, and minor allele frequency of 5%. In the analysis, only biallelic SNPs are included. D statistic between all our trios was calculated with the phylogeny as a reference tree to guide the analysis using the Dtrios tool in the Dsuite toolkit (v.0.4). This compares all possible trios of species and calculates a genome-wide minimum D, as well as a significance value of each trial, it will also calculate F4 statistics for each trio that we used to calculate Fbranch statistic that we used to infer signals of past introgression. We used the DInvestigate tool from Dsuite to generate window-based measures of f_dM, as a measure of introgression for each trio that showed significant (BH corrected p < 0.05) introgression in the genome-wide analysis (Fig. 1 e) in overlapping windows of 50 SNPs (25SNP overlap). We were interested to see if there were any trios that showed an elevated signal of introgression around the BarH1 locus. To this end, we filtered out all trios that showed an increase in f_dM in the 600Kb region surrounding BarH1, this revealed 3 species trios (Supplementary figure 3). However, when investigated in detail, they were found to represent increased haplotype similarity between 2 fixed alba species in comparison to fixed orange species; additionally, none of these had any increase in f_dM across the actual BarH1 genic region or across the insertion region.

#### GWAS of Alba in C. eurytheme

DNA was extracted using KingFisher Cell and Tissue DNA Kit from ThermoFisher Scientific (N11997) and the robotic Kingfisher Duo Prime purification system. DNA purity was assessed using 260/280 ratio (Nanodrop 8000 spectrophotometer; Thermo Scientific, MA, USA) and concentration was quantified on a Qubit 2.0 Fluorometer (dsDNA BR; Invitrogen, Carlsbad, CA, USA). DNA was sent to the Science for Life Laboratory (Stockholm, Sweden) for library preparation and sequencing (Rubicon, 150bp PE reads with 350bp insert size, Illumina HiSeqX). Sequencing libraries were sequenced twice, to a predicted depth of 10x each time. Raw reads were clone filtered, had adaptors trimmed and low-quality bases (PHRED 20) removed using the BBduk tool from the BBmap software package v34.86 (Bushnell B. sourceforge.net/projects/bbmap/). Cleaned reads were mapped to the C. eurytheme reference genome using NextGenMap v0.5.2. SAMTOOLS was used to filter out unmapped reads, sort, duplicate marking of reads and indexing. PICARD-tools v1.139 was used to add Readgroups and then again to merge the separate sequencing runs of each sample, before a final round of duplicate marking using SAMTOOLS. Variants were called using Freebayes v1.3.1-16-g85d7bfc (Garrison E −2012 arXiv), and the resulting VCF-file was filtered using a mix of VCFTOOLS v0.1.13 Danecek, (Petr, et al. - 2011) and custom awk scripts. The final GWAS was run using PLINK v1.9 (Chang et al. - 2015) to identify associated loci. We filtered the GWAS based on two separate sets of filters were used. First, we filtered the VCF file to remove low quality SNPs or sites that did not match our depth or frequency criteria (minimum depth 3, minQ 20, max-missing 0.95, and minor allele cutoff of 0.05). Second, we added an additional filter to this set added the prior criteria to include only sites that were unique to the Alba females. Additionally, we also filtered the output using the information gained from the QTL analysis, and only kept SNPs on the scaffolds that make up Chromosome 3.

The same filtering approach was also applied to the GWAS done using the Alba reference genome.

#### Generation of draft genomes alba individuals of C. eurytheme, C. nastes and C. crocea

DNA from wild caught females was extracted from the thorax of samples stored frozen in 95% ethanol using the same protocol as in the resequencing done for the GWAS. Prior to library preparation an additional analysis of molecular wight of the DNA was performed using gel electrophoresis (0.5% agarose LE). DNA was extracted from 2 individuals of each species and the ones with the highest quality DNA, as determined by gel electrophoresis, nanodrop measurement and Qbit were selected for library prep. Sequencing and assembly with Supernova v2.1.1 at SciLifeLab.

#### PCR-based validation of insertion

The presence and uniqueness of the insertion to Alba individuals were further validated using PCR-based markers. The primers were designed to bind within the insertion region. that was found to be unique to Alba. Optimal primer binding sites were identified using the primer3 software (libprimer3 release 2.5.0, Untergasser et al. - 2012). PCR-reactions were run on DNA extracted from 8 orange and 8 Alba females, that had not yet been sequenced or used in the GWAS. Positive controls were run using previously validated primers binding to mitochondrial cytochrome oxidase I gene (Brower et al. 2006). The reactions were run using Invitrogen Platinum Taq in a Veriti 96-well thermocycler (Applied Biosystems, Foster City, CA, USA) using the recommended settings for the polymerase (72C x 2min followed 35 cycles of 94C x 30sec + 54C x 30sec + 72C x 15 sec followed by 72C x 5min). The PCR product was visualized by agarose gel electrophoresis in a 1% gel (Supplementary figure 7).

#### Generation of Alba reference genome

We identified the scaffold containing *BarH1* in the supernova assembly of the *C. eurytheme* using tBLASTn. We then aligned all the resequencing data from the GWAS to this contig alone and evaluated along the contig visually in IGV. Regions where no orange reads aligned, but Alba did, were extracted and blasted back against the *C. crocea* reference genome (Woronik 2019) to assess whether this was the previously identified Alba insertion region. Once we had established the Contig containing both BarH1 and the Alba insertion, we blasted the entire contig back against the Orange *C. eurytheme* reference genome to establish the edges of the contig in the reference genome as well as to establish orthology. Once edges were established, we manually inserted the entire contig, making sure to remove the overlapping sequence, generating what we refer to as the Alba reference genome.

We decided to generate the additional *C. crocea* assembly to make sure that we would be able to assemble and subsequently detect the alba insertion sequence. This was done using the same method as described in the main paper for the detection of the Alba allele in the *C. nastes*, and *C. eurytheme* assemblies, where we blasted the previously known alba allele detected in the *C. crocea* assembly from Woronik et al. 2019, and if this insertion was co-located with the BarH1 gene. We also assessed sequence similarity and synteny using Blastn, which we visualized using Kablammo (supplementary figure 8 & 9).

#### Alignment and assessment of the Alba insertion across species

The sequence data generated for the phylogeny was aligned to the Alba reference-genome using NextGenMap, filtered for mapQ 20 and being in proper pairs. Mapability of the data, and confirmation of sex in cases where it was uncertain was done using read coverage analysis using goleft indexcov (see SM_CovStats). This confirmed that 7 samples in our analysis were indeed male (Colias erate poligraphus, Colias tam. mogola, Colias phicomone, Colias Tyche, Colias behri(B033), Colias tamerlana (B037) and Colias wiscotti) based on the increased coverage on the scaffolds that make up the sex-chromosome (9, 19 & 52). It also emphasizes the divergence between the South American species *C. lesbia* and *C. euxanthe*, as those species consistently showed reduced coverage on the terminal ends of the shorted chromosomes in the assembly (SM_CovStats).

Coverage across the insertion region was then visually inspected visually in IGV. We especially assessed the regions where we had observed a difference in coverage between Orange and Alba eurytheme individuals and whether this extended to other species where we had sampled either Orange or Alba individuals. The sequence that was unique to only white-colored species (putative Alba species) was the conserved Alba region and likely causal for the phenotype across Colias.

To establish a null expectation for how often a region of the same size as the conserved Alba region would segregate in this manner between white and colored species (e. g orange; yellow; red) we performed a read depth analysis of all the sequenced *Colias* species. Reads were aligned using Nextgenmapper and filtered for being good pairs. We then used goleft v.0.2.1 (https://github.com/brentp/goleft) to evaluate the average read depth of 600bp windows across all scaffolds. We selected 600 basepairs rather than the full 1200bp of the Candidate alba locus to ensure that at least one window ended up inside the insertion region, this also made the analysis more sensitive compared to using a larger window size. Read depth in the windows was then binary classified as either having read coverage or not, with the latter classification assigned if they had less than 25% of the read depth compared to the scaffold average. Thus, for each window, each species had a value of 0 or 1. Then, using these values of either having reads covering the window or not, we calculated the mean value per window for the white and colored groups of species, then calculated the white minus colored value. For the Alba identified region in white-colored butterflies, this value was 1, which was then compared to the rest of the windows across the genome, which served as a genome scale control.

#### Phylogenetic analysis of the Alba insertion

We extracted the consensus sequence of the region from all 43 Alba samples in our analysis (14 *eurytheme*, ten *crocea*, seven *philodice* (Maryland), 1 philodice (from British Columbia) and one from each remaining alba species in the phylogeny) using the bam2fasta 1.4 tool-kit. We keept all sites with at least one read covering it and selected the most common allele at polymorphic sites. We then ran IQtree to generate a phylogenetic tree of the sequences. IQtree was run with model finder plus enabled to allow it to find the most parsimonious model.

#### CRISPR/Cas9 targeted mutagenesis of the Alba insertion

We used the PROMO tool to narrow down regions of interest in the Conserved Alba locus, while we did keep an out for candidate genes that have been previously shown to either interact with BarH1 or known to be involved in sexual dimorphisms, such as *Doublesex*, we also used it to get an idea of sites that showed a higher density of many different potential transcription factors. In the end, we selected four target sites, two that targeted the only putative binding site for the *Doublesex* transcription factor and two that targeted upstream and downstream of this position. Together, if all cuts would be successful, this would remove ~500basepairs of sequence and remove a large portion of the potential binding sites. We did four types of cocktails of gRNA/Cas9 mixture that we injected;

gRNA_P(doublesex binding site)
gRNA_P and gRNA_U(3bp upstream of doublesex binding site)
gRNA_2 (~250bp upstream of gRNA1) and gRNA_5 (~150 bp downstream of gRNA1).

And all four together.

Unfortunately, due to poor egg-laying of the alba females, we could not inject an equal number of eggs with each combination, instead primarily injecting eggs with either gRNA_P alone, or gRNA_U and gRNA_P.

The only combination we saw any phenotype from was when we injected all 4 together. But as this was only done once the females had almost stopped to inject eggs, we didn’t inject more than 40 eggs with this combination. The major cause of death among our injections was the larvae getting stuck in the double-sided tape that was used to fixate the eggs to a glass slide during injection. Out of 200 injected eggs, we were only able to transfer 39 larvae to fresh hostplant (5 of the 40 eggs we injected with all four gRNAs).

#### Validation of CRISPR/Cas9 targeted mutagenesis

Using Primer3 we designed PCR primer pairs that bind upstream and downstream of gRNA_2 and gRNA_5. Due to a large amount of repetitive DNA in the region, we were forced to have the forward primer bind to a non-unique location, leading to a risk of having some off-target bind sites. The reverse primer was unique, though, and there were no alternative bind-sites for the forward primer within 200Kb of the intended location. The reactions were run using Invitrogen Platinum Taq in a Veriti 96-well thermocycler (Applied Biosystems, Foster City, CA, USA) using the recommended settings for the polymerase (72C x 2min followed 35 cycles of 94C x 30sec + 54C x 30sec + 72C x 15 sec followed by 72C x 5min). The PCR product was visualized by agarose gel electrophoresis in a 1% gel (Supplementary figure 12). We used the primer pair developed by Woronik. Et al 2019. to validate the Alba status of the sample and as a positive control for the reaction, this primer pair does not bind in the vicinity of each other.

### Tables

**Stable1.**
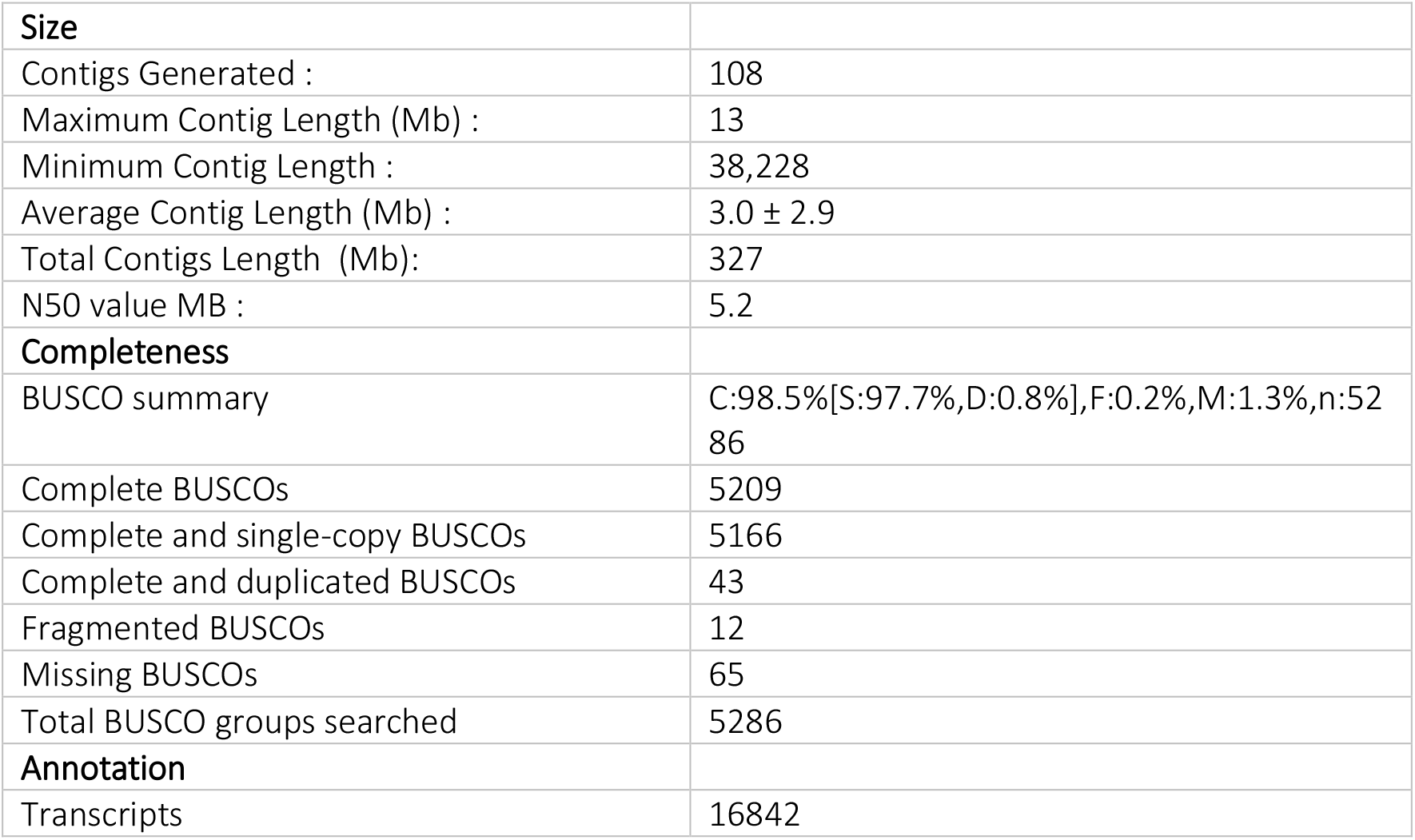
Quality metrics of the assembly

ST1. Metadata table of sequenced individuals, including capture site, sequencing depth, and sample name.

ST2. Top hits of GWAS after light filtering of SNPs, using the orange reference.

ST3. Top hits using informed priors, using orange reference

ST4. Top hits using QTL informed priors, orange reference

ST5. Top hits using Alba reference genome and light filters. Note that the hits not on the alba locus are on the sex chromosome.

ST6. gRNA sequence

ST7. Injection statistics and survival.

ST8. PCR primers.

ST_covstats. Normalized coverage of reads across scaffolds of all samples.

### Figures

**Supplementary flowchart 1.**
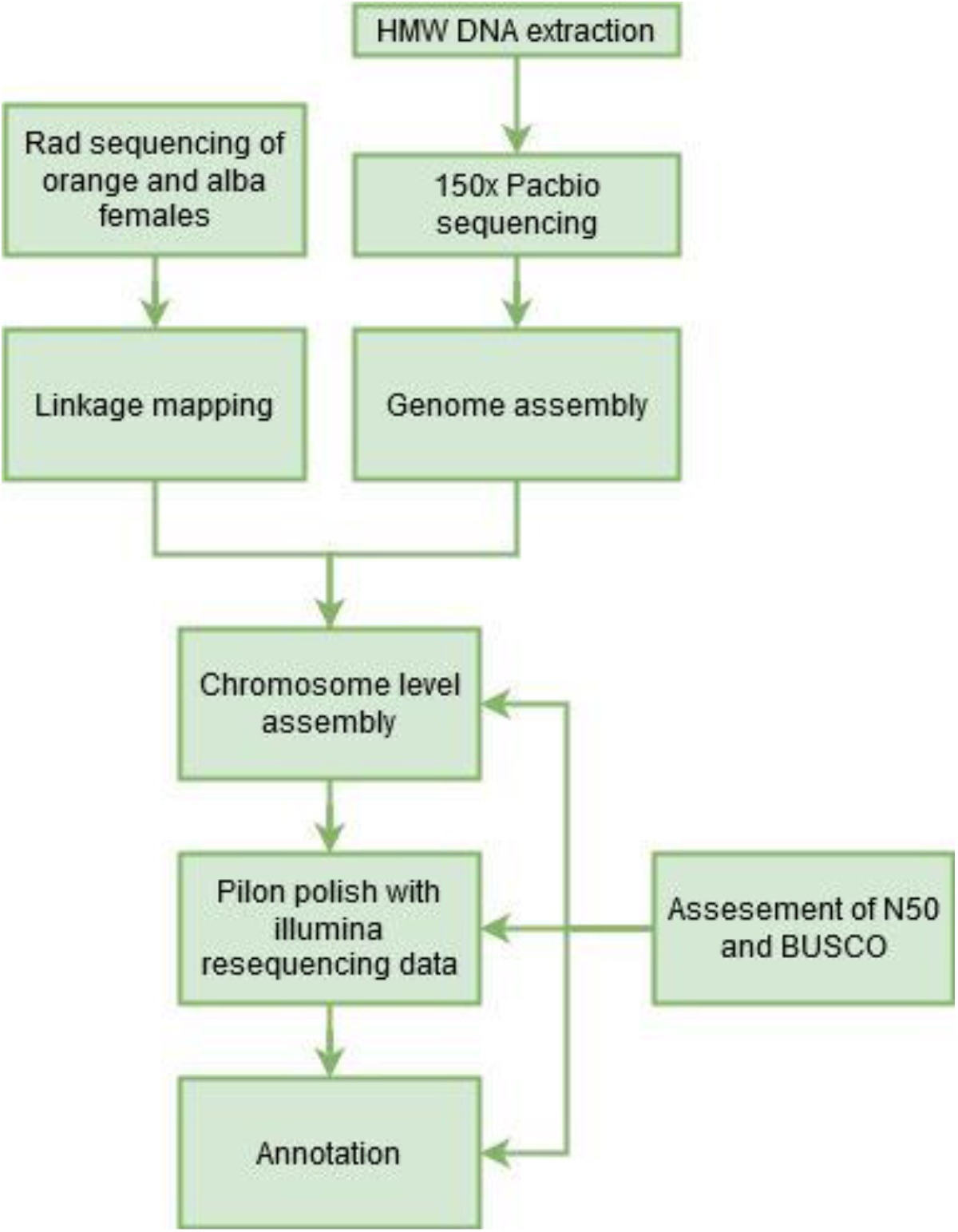
Assembly pipeline. Flowchart illustrating the genome assembly, annotation, and evaluation process. Generation of raw contigs was done using 150x Pacbio sequence data coming from a single female pupa. Linkage mapping of the contig data was done using Rad sequencing from multiple individuals of *C. eurytheme X C. philodice* hybrid crosses. The draft assembly was then polished using Pilon to reduce indel errors introduced by the pacbio sequencing, and finally annotated using BRAKER2. Each step of the assembly process was analyzed using N50 and BUSCO.

**Supplementary flowchart 2.**
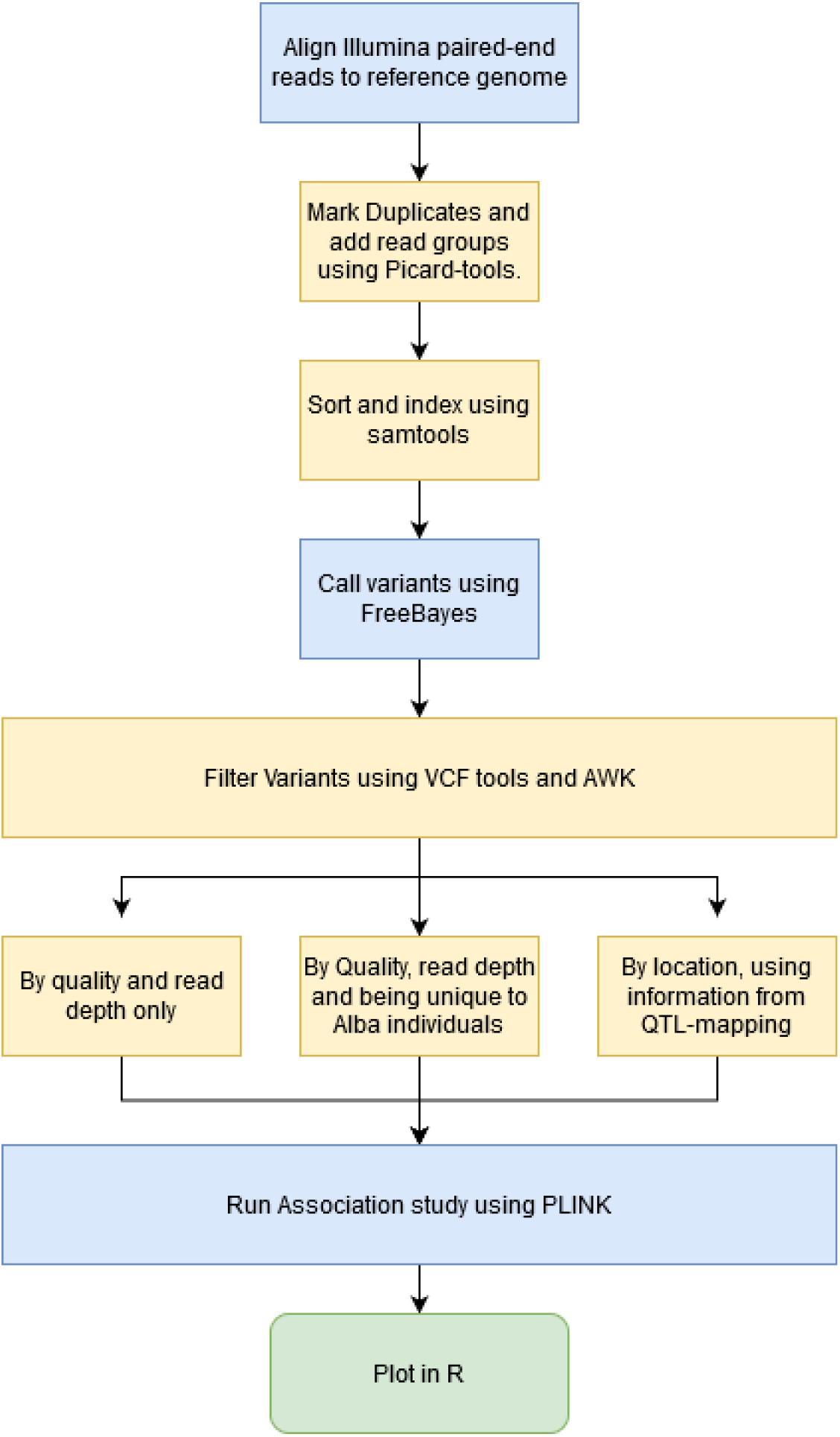
Variant calling pipeline. Flowchart describing the steps involved in the Genome Wide Association study (GWAS). Blue boxed represent the generation of Data, Yellow boxed filtering of data and green boxes visualization. In total we ran 3 levels of filtering of the raw VCF-file with different level of priors.

**Supplementary figure 1.**
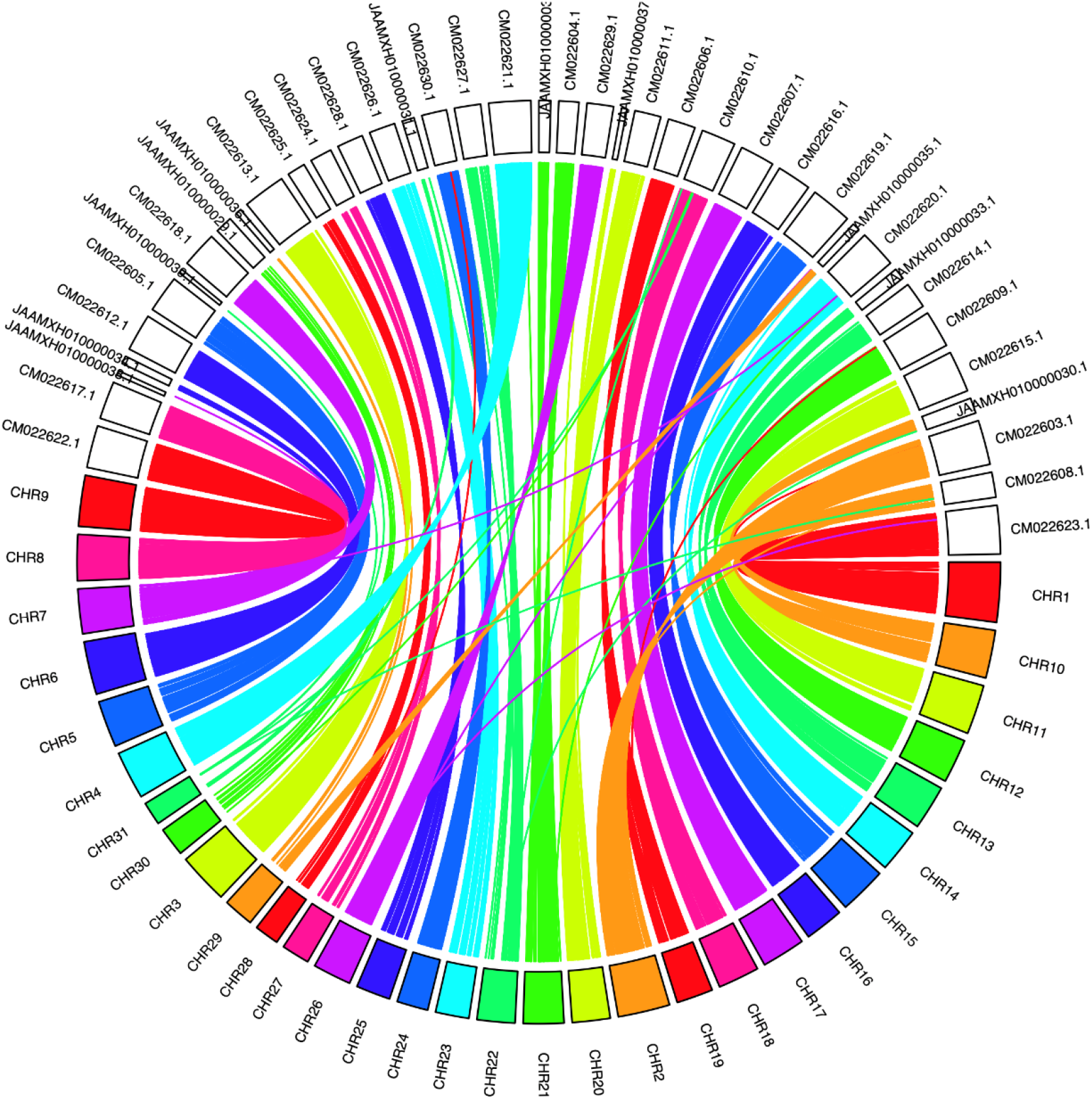
Syntheny plot illustrating the conserved chromosomal structural comparison between the Colias eurytheme genome and Zerene cesonia. Shown are the 31 linage groups identified in C. eurytheme colored by chromosome, while the scaffolds of Z. cesonia are uncolored. Each line represents an inferred orthologous region. Only scaffolds > 1 Mb of *Z. cesonia* were included, with nucleotide alignment identity > 88 %. Note how several Z. cesonia scaffolds are brought together within C. eurytheme chromosomes (e.g. Chr 6 and Chr 29). Given the high synteny between *Z. cesonia* and *Heliconius erato* (Rodriguez-Caro et al. 2020), and *Heliconius* to other butterflies and moths (Ahola et al. 2014), we infer that *Colias* chromosomal structure adheres to the standard Lepidoptera chromosome structure (Hill et al. 2019).

**Supplemental figure 2.**
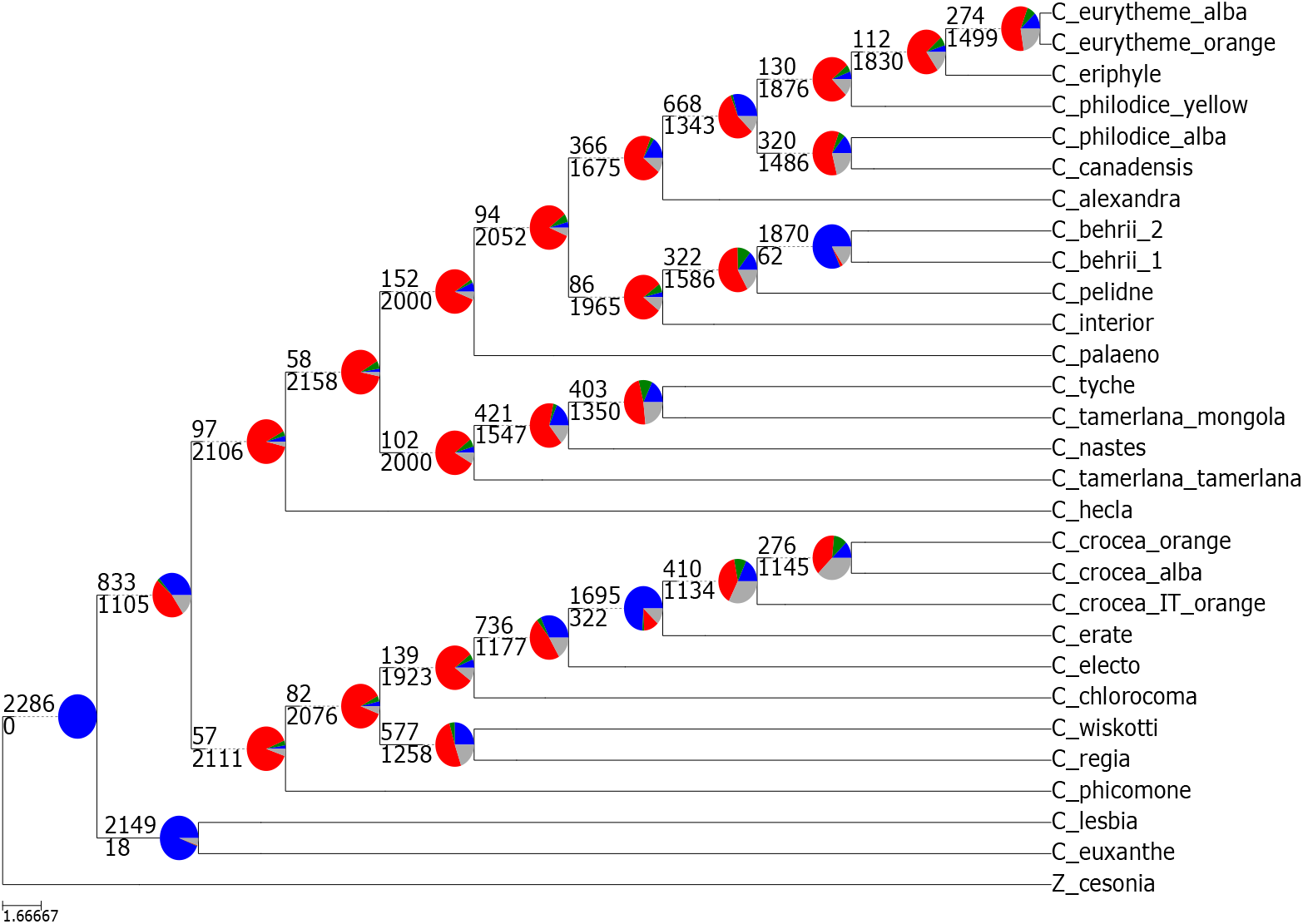
Pie charts of gene tree concordance, conflict, and lack of signal compared to the Astral species tree. While traditional node support values, whether bootstraps or posterior probability, generally overstate gene tree support, here we present direct quantification per node of gene tree topology with the species tree topology. At each node, the number of gene trees concordant (top number) and in conflict (bottom number) is shown. Pie charts at each node give a further breakdown of gene tree proportions by those concordant with species tree (blue), those that support a common alternative topology (green), those that support the remaining low-frequency alternatives (red), and those lacking robust information as the have less than 50% bootstrap support (gray). Here, the South American taxa are very well supported, as are the clades containing: *C. crocea, C. erate, C. electo; C. tyche, C. tamerlana mongola, C. nastes; C. pelidne, C. behrii; C. alexandra, C. canadensis, C. philodice, C. eurytheme, C. eriphyle*. Note how the vast majority of nodes are red, indicating extensive gene tree conflict rather than lack of phylogenetic signal (gray).

**Supplementary figure 3.**
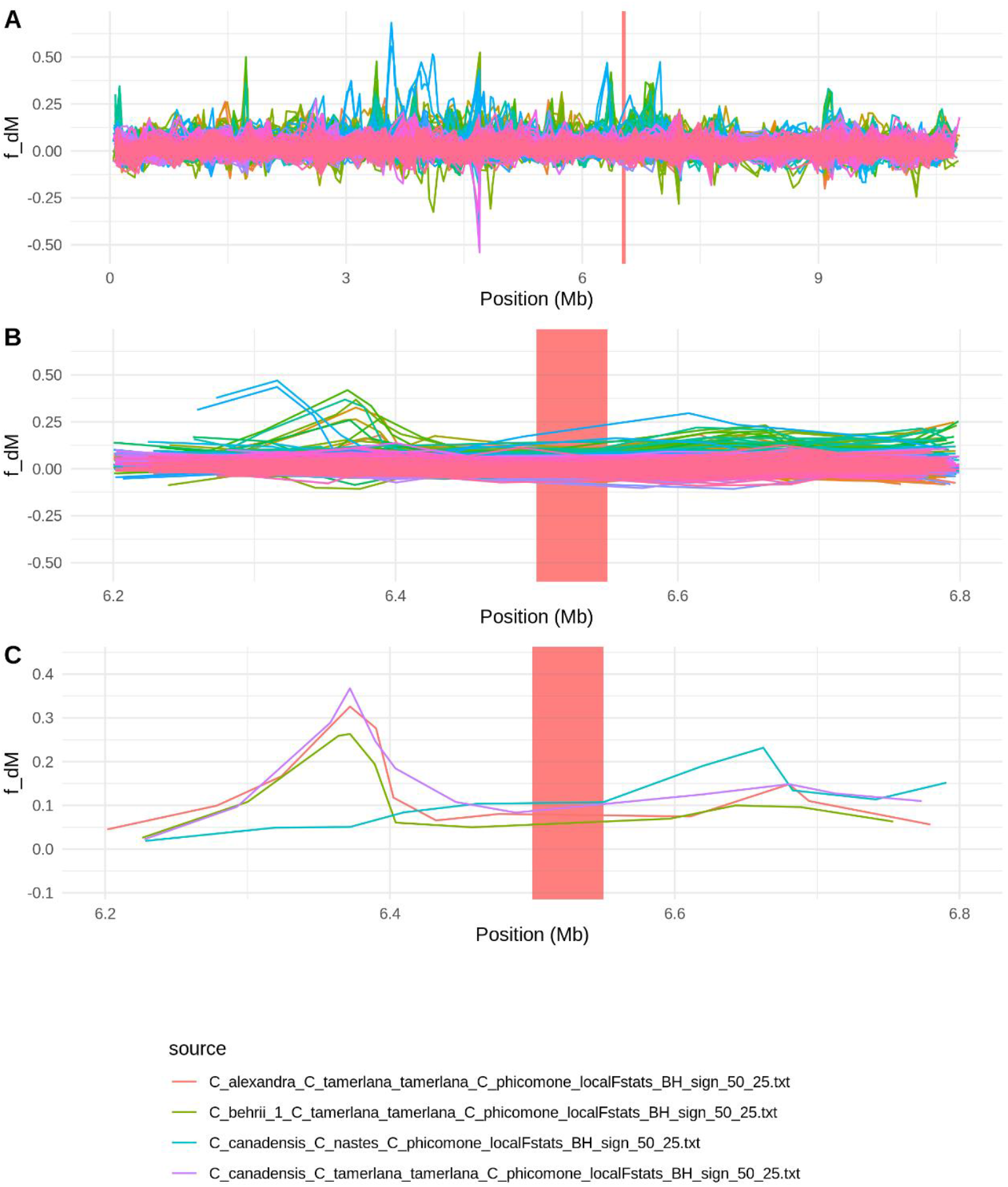
Assessment of signals of localized introgression. By looking at f_dM in sliding windows of 50 SNPs across the genome for all species trios showing significant levels of introgression in the genome-wide Introgression analysis looking at A. Whole scaffold containing BarH1, B. 600Kb region surrounding BarH1 and C. The four species trios with an f_dM higher than 0.2 anywhere in the 600Kb region surrounding BarH1. The BarH1 region is highlighted red in all three plots.

**Supplementary figure 4.**
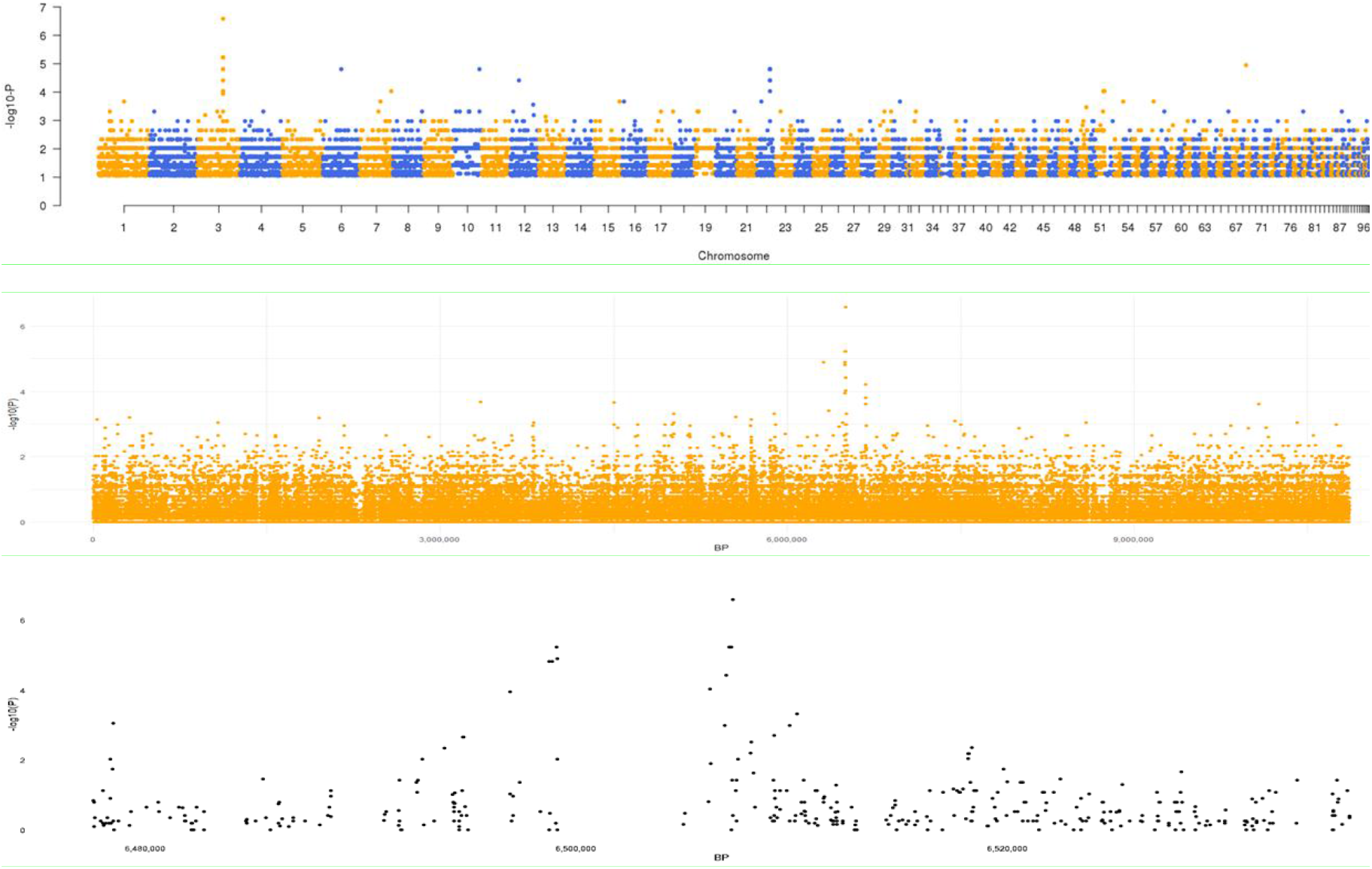
GWAS results. GWAS against the Orange reference genome using the light filters of quality and depth only. -log10(p-value) for SNP correlating with alba color against the position in the genome. Panel A: Genome-wide, B: Sc0000002, C: 60kb around the highest peak.

**Supplementary figure 5.**
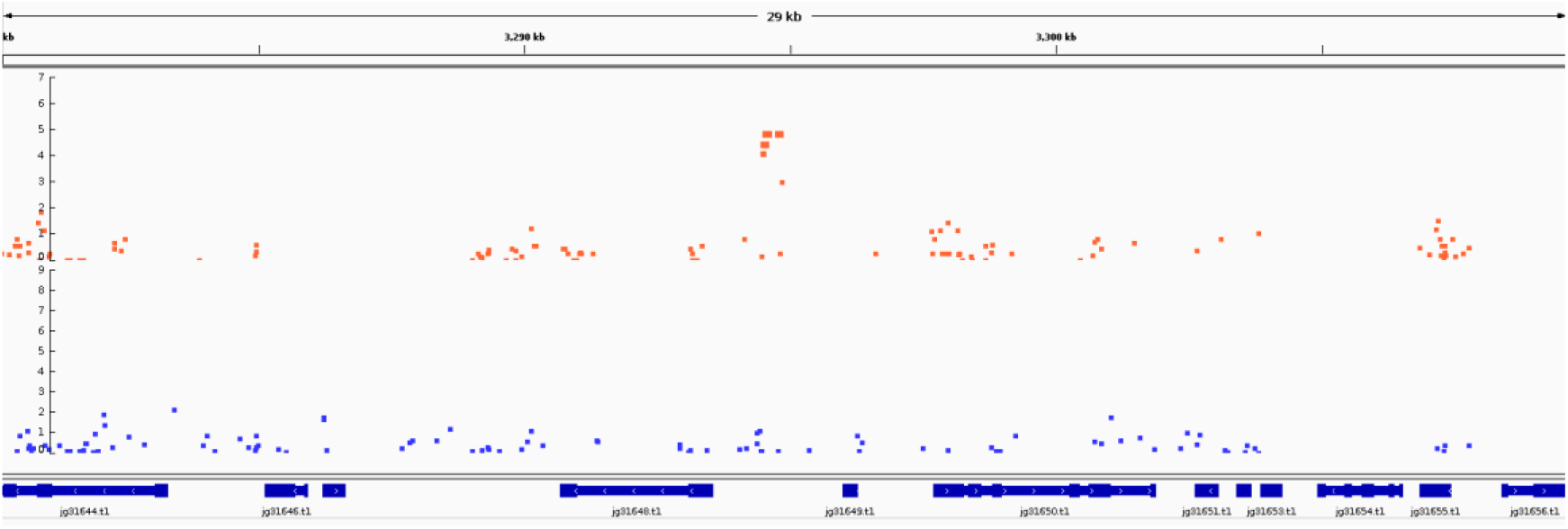
Close up on the second GWAS locus identified on Scaffold 22 when the orange reference genome was used. The top row colored in orange represents sites identified against the orange reference genome, while the blue are from the synthetic Alba reference genome. The gene to the right of the highly associated loci is similar PIFI-like transposase when blasted against NCBI, and the gene to the right is similar to a PiggyBac transposon.

**Supplementary figure 6.**
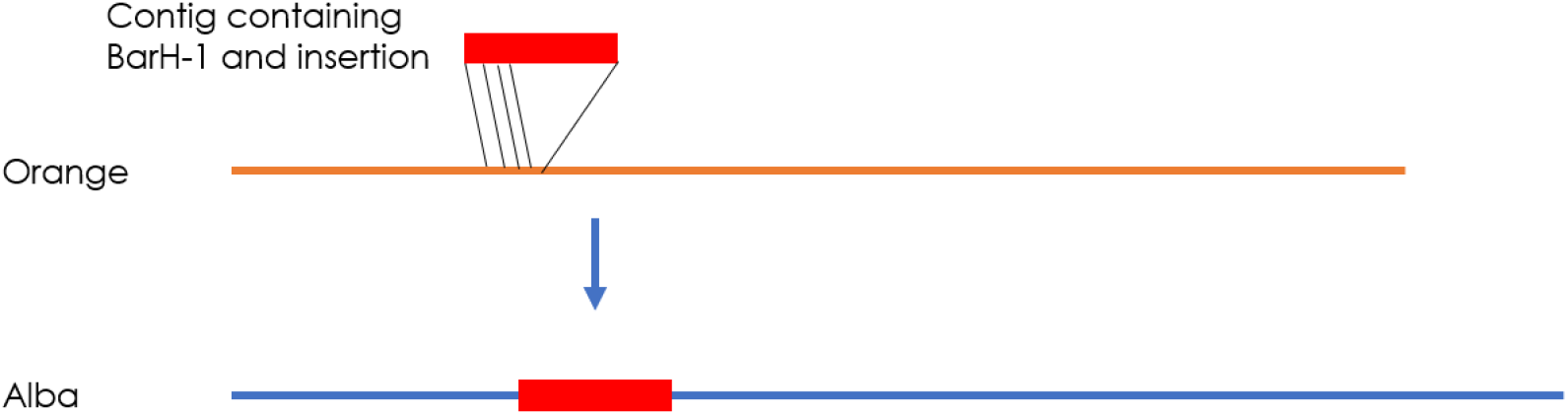
Illustration of the generation of the Alba reference genome. The contig containing *BarH1* and the insertion was identified using blast and read depth analysis. Overlapping sequences between the contig and the orange reference genome was used to define the edges at which to insert the sequence. The sequence was then inserted and overlapping sequences removed, favoring the Alba Contig sequence leading to the generation of the Alba reference genome.

**Supplementary figure 7.**
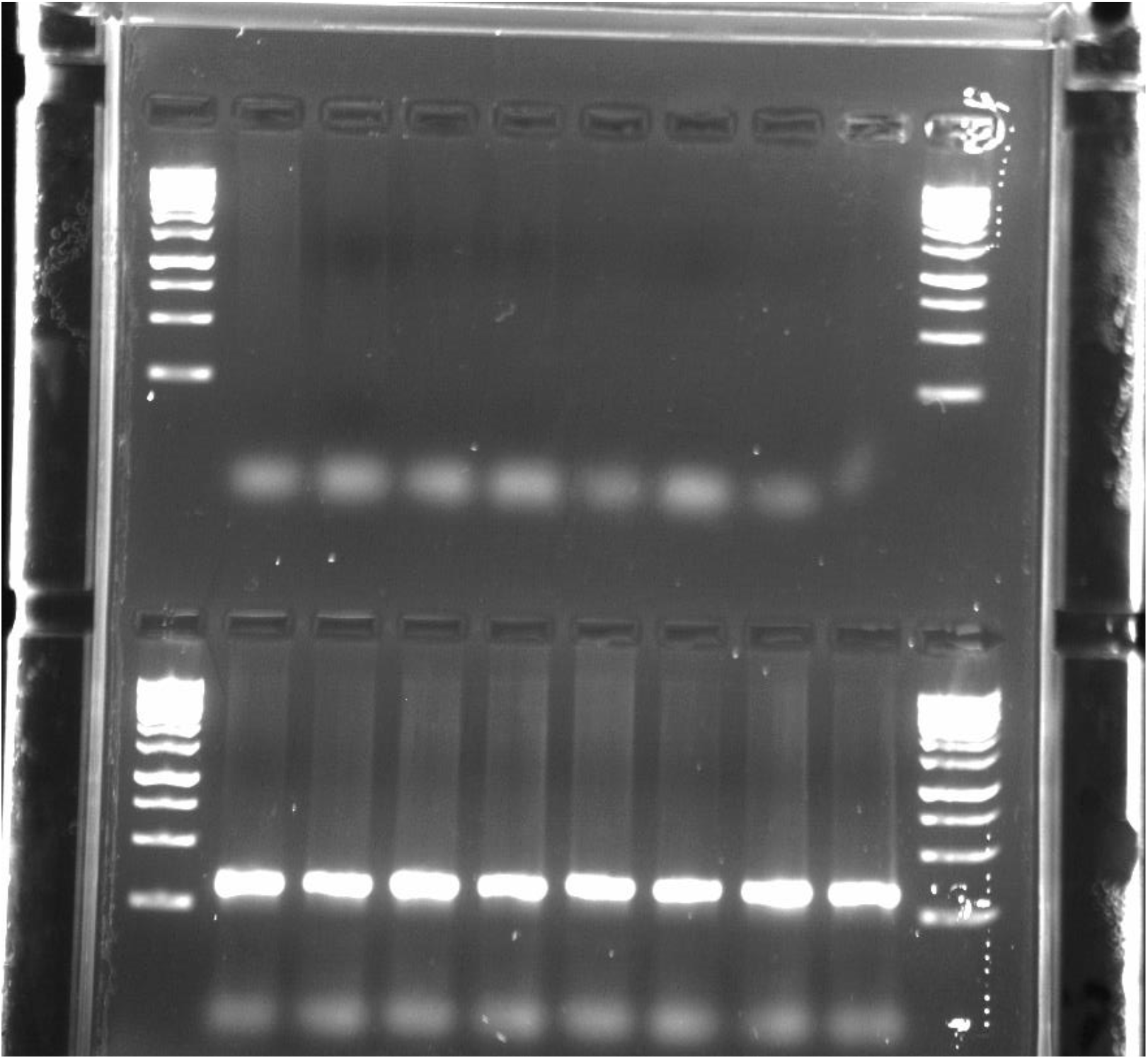
Verification of the Alba locus using PCR-based markers. 8 orange and 8 Alba *C. eurytheme* individuals, independent from the GWAS analysis, had DNA extracted and then genotyped for the insertion. PCR products were visualized on a 1% agarose gel. The top row shows negative results in orange females, and the bottom row shows positive results from alba females.

**Supplementary figure 8.**
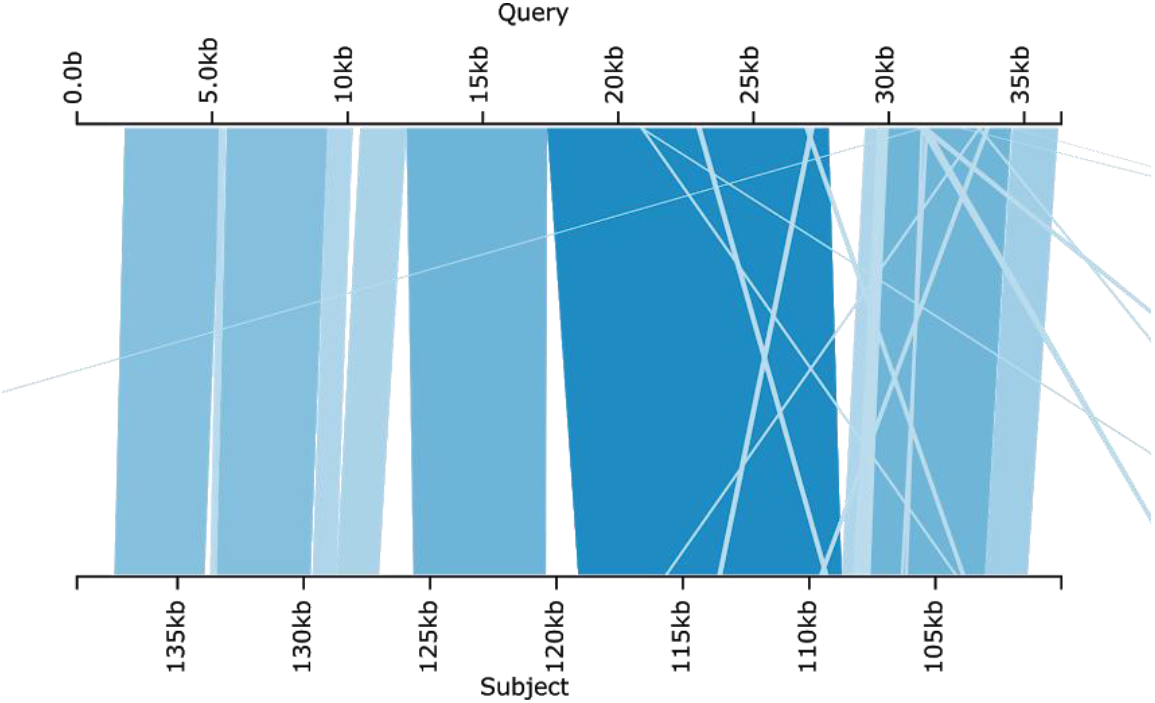
Orthology assessment of the C. crocea alba scaffold identified in the Chromium 10X assembly. (query), identified by blasting the C. eurytheme BarH1 gene and insertion sequence against the assembly, compared to the one found in the C. crocea reference genome(subject). Darker blue hue means higher similarity.

**Supplementary figure 9.**
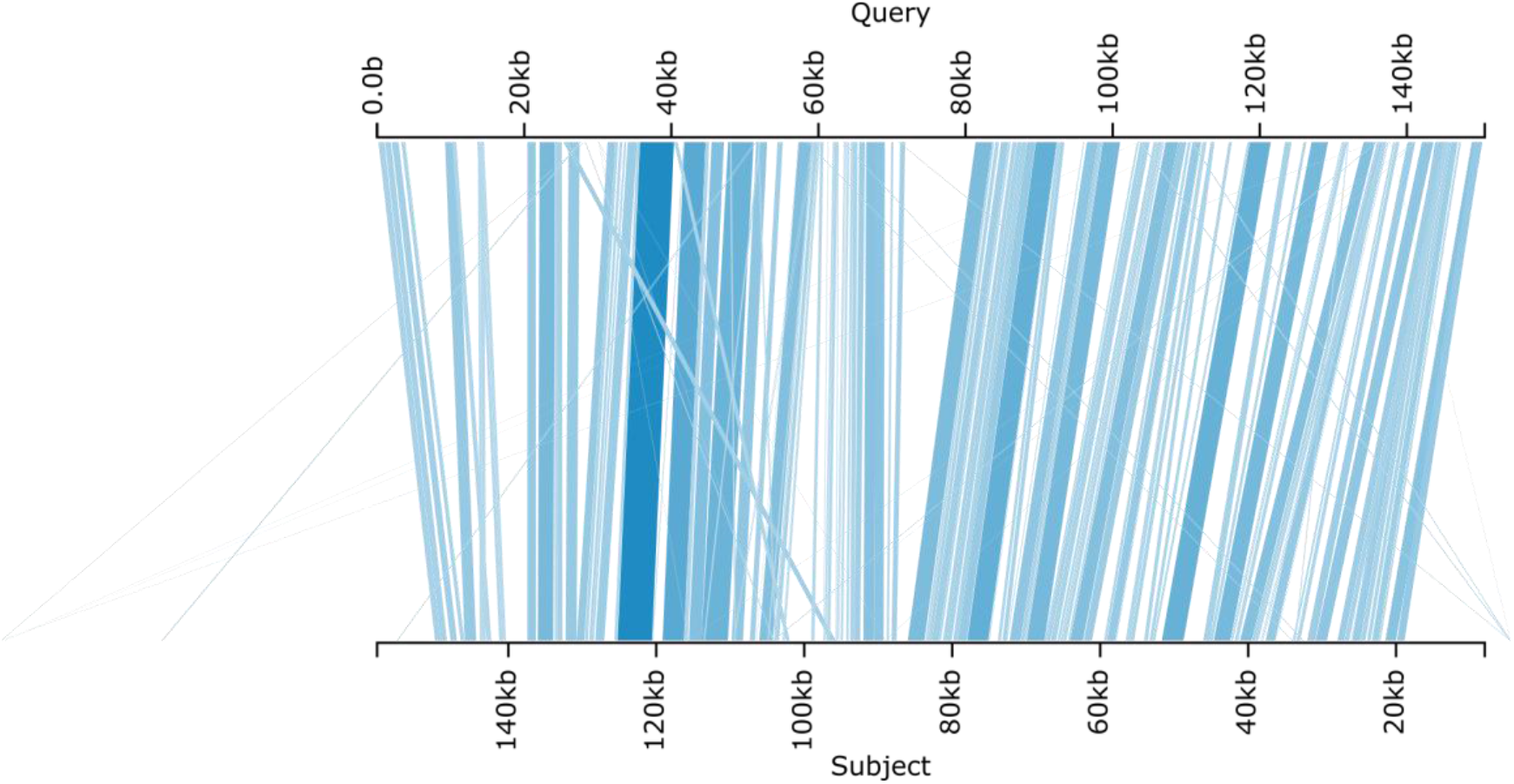
*Orthology assessment of the Alba insertion in* C. nastes. The alba scaffold, as identified using the *BarH1* gene and insertion identified in *C. eurytheme*, from the *C. nastes* Chromium 10X assembly(*query*), aligned against the Colias crocea reference genome(subject). Darker blue hue means higher similarity

**Supplementary figure 10.**
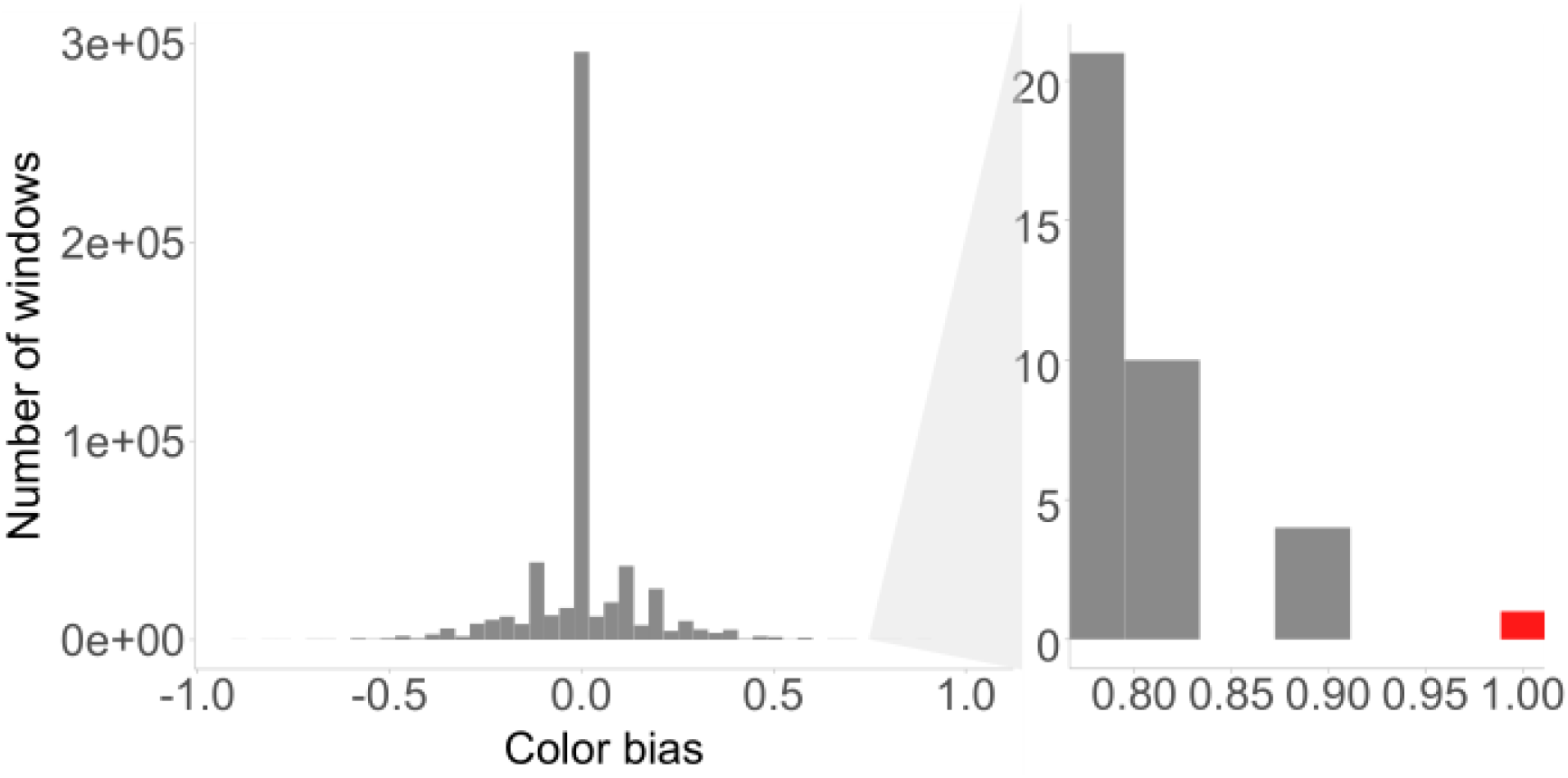
Read depth analysis between Orange and Alba species in 600bp windows. Read depth was calculated for each species in 600bp windows when the reads were aligned against the synthetic alba reference genome. Each window was then classified as either having reads covering or not if the read coverage was less or more than 25% of the mean scaffold depth for the species. A mean was then calculated for all the alba species and all the Orange species, and finally we subtracted the orange mean from the Alba beam. Only 1 window, the conserved alba locus, had zero coverage in the Orange individuals, and complete coverage in the Alba. No window showed the opposite pattern.

**Supplementary figure 11.**
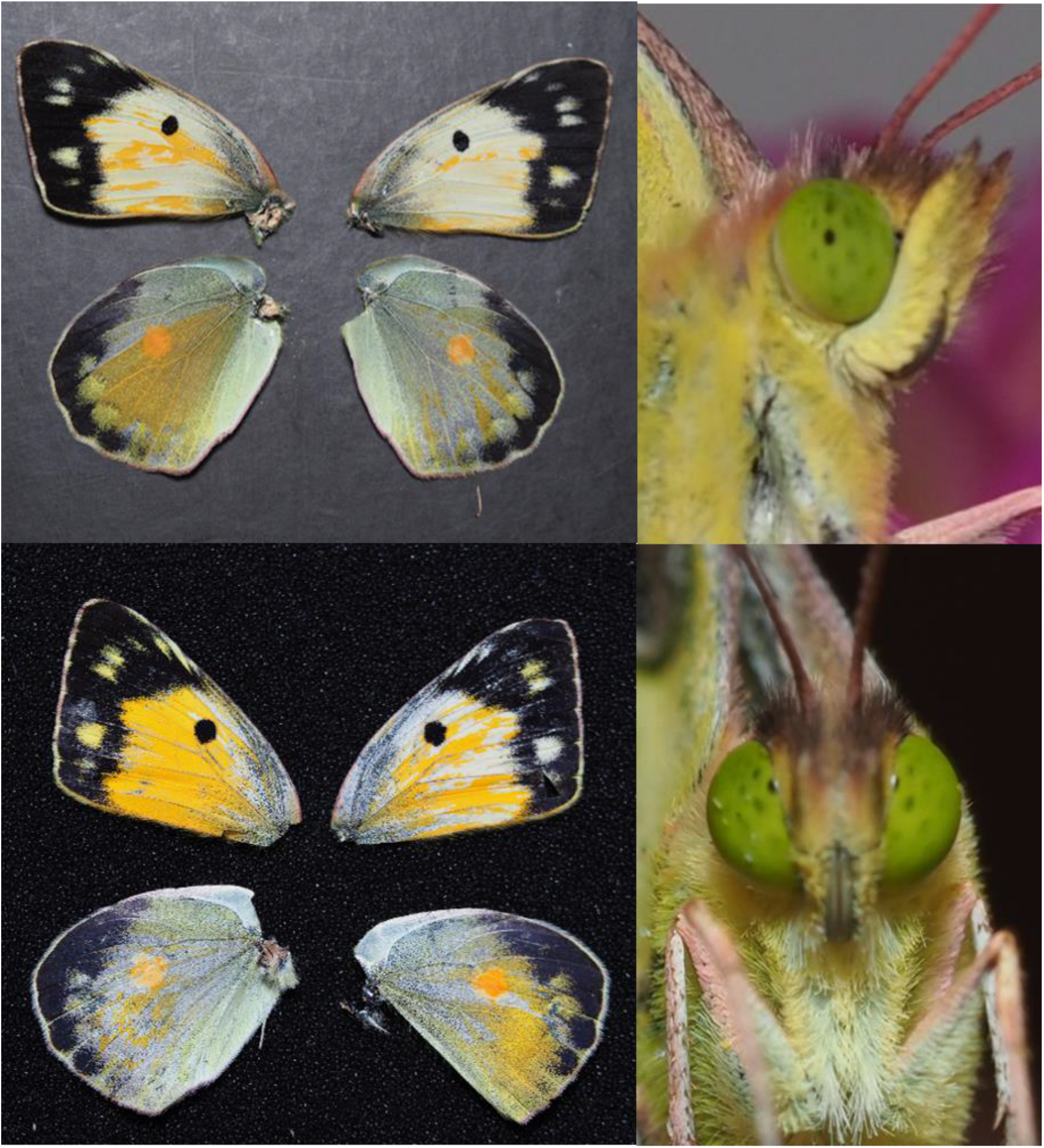
Alba CRE-KO. Images of both successful “conserved Alba region” CRISPR-KOs with their phenotypes: ind. Alba CRE-KO 1 wing (top left), ind. Alba CRE-KO 2 wing (bottom left), eye ind. Alba CRE-KO 1 (top right), eye ind. Alba CRE-KO 2 (bottom right)

**Supplementary figure 12.**
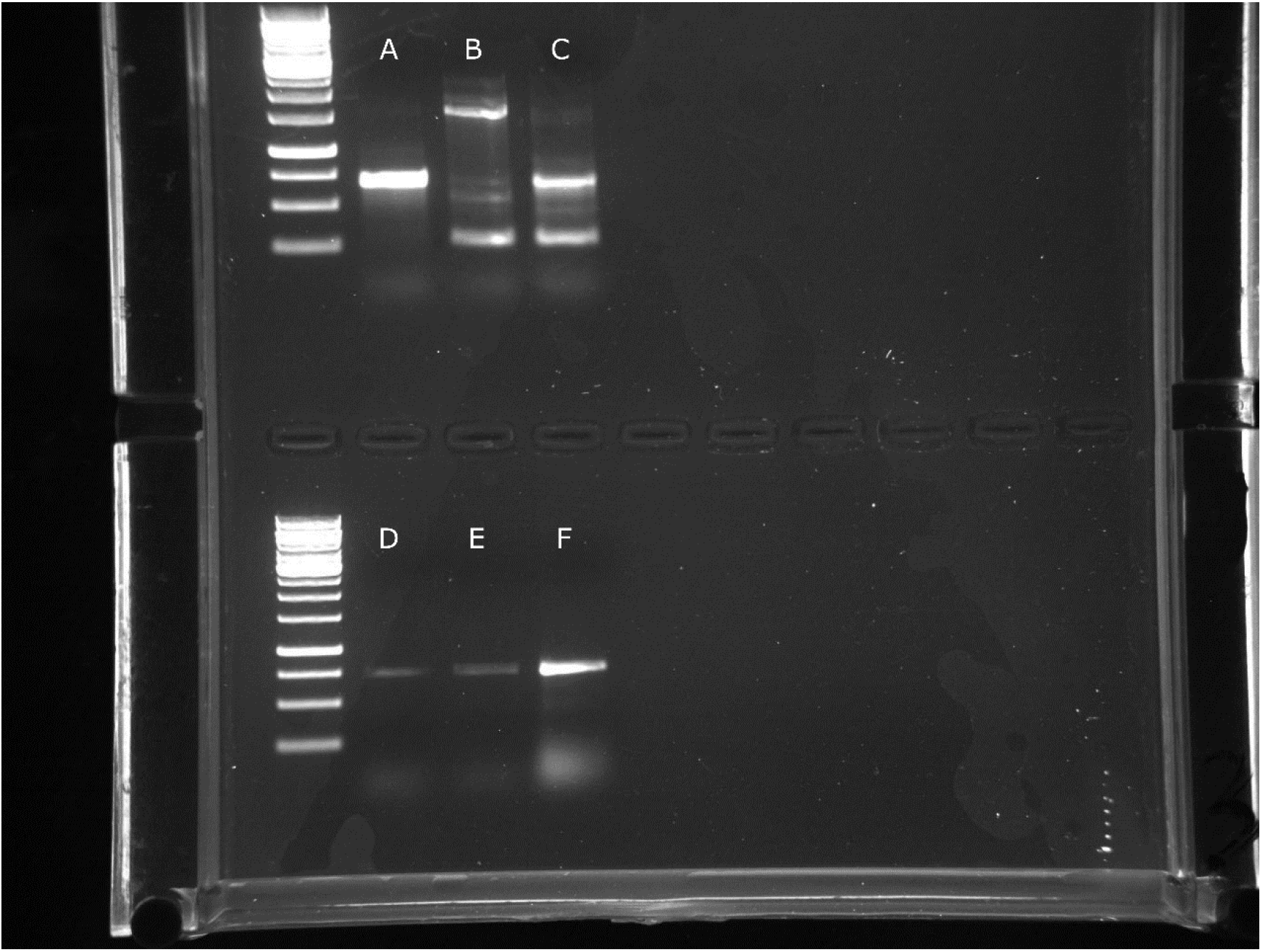
PCR validation of CRISPR-induced deletions. Gel verifying CRSPR KO as well as alba status for the two successful mutants. From left to right the well show, **A**: Alba-CRE WT, **B**: Alba-CRE KO-1, **C.** Alba-CRE KO-2, **D:F.** Alba control validation primer on the sample in the column above. Not the variation in band sizes caused by the Cas9 in the two knockout samples.

**Supplementary figure 13.**
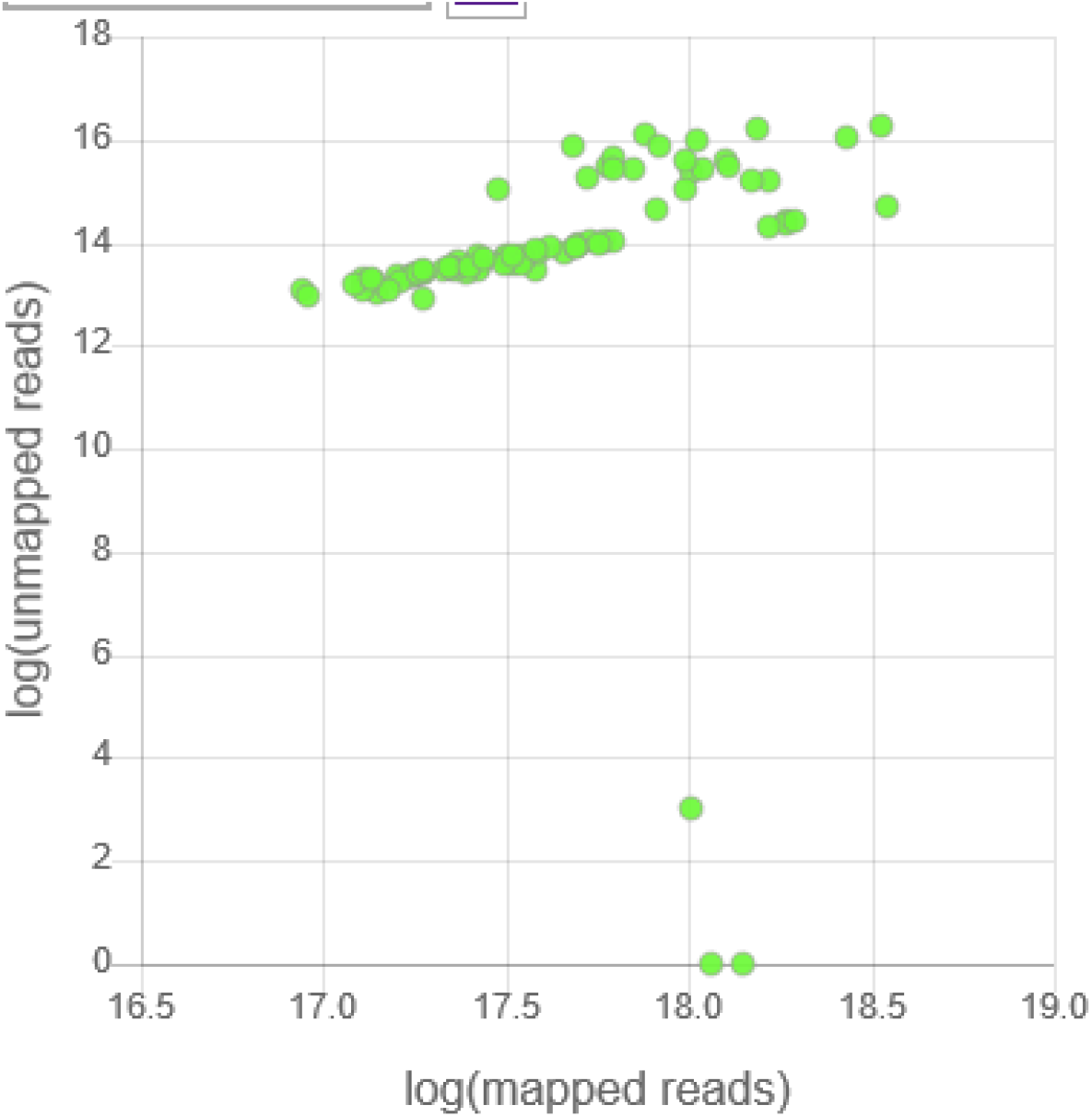
Mappability of short read datasets. Scatterplot showing counts of mapped reads to unmapped reads. Note that the samples with very few unmapped reads had been filtered for mapped reads prior to the generation of this figure.

